# Biosynthesis of oxyresveratrol in mulberry (*Morus alba* L.) is mediated by a group of p-coumaroyl-CoA 2’-hydroxylases acting upstream of stilbene synthases

**DOI:** 10.1101/2024.04.04.588114

**Authors:** Antonio Santiago, Pablo Romero, Ascensión Martínez, María José Martínez, Jone Echeverría, Susana Selles, Raquel Alvarez-Urdiola, Chen Zhang, David Navarro-Payá, Gastón A. Pizzio, Estel·la Micó, Antonio Samper, Jaime Morante, Riccardo Aiese Cigliano, David Manzano, Roque Bru, José Tomás Matus

## Abstract

Mulberry (*Morus alba* L.) is considered a millenary medicinal plant and a food source for silkworms. Different *M. alba* extracts offer a variety of biological and pharmacological properties that are in part attributed to stilbenoids, a small group of phenylpropanoids that include resveratrol and oxyresveratrol. These are naturally present in non-renewable parts of mulberry trees, impeding their efficient extraction. As a way to bypass this spatiotemporal restriction, we generated cell suspensions from mulberry twigs and demonstrated that the combined use of methyl jasmonate and methyl- or hydroxypropyl-β-cyclodextrins elicited a high production of resveratrol and oxyresveratrol, both intra and extracellularly. To identify oxyresveratrol-producing enzymes (unknown to date), we first improved the structural and functional annotation of the mulberry genome by integrating short and long-read sequencing data. We further combined this data with transcriptome, metabolite and proteome time-series evidence to identify a complete set of elicited phenylpropanoid- and stilbenoid-related genes. These included 22 stilbene synthase (*STS*) genes and a group of six *p*-coumaroyl-CoA 2’-hydroxylases (C2’Hs) that were highly co-expressed with resveratrol and oxyresveratrol accumulation. We transiently transformed *Nicotiana benthamiana* plants and grapevine (*Vitis vinifera* L.) cell suspensions to functionally validate the role of C2’Hs as the first committed step of oxyresveratrol synthesis, providing an alternative substrate for STSs by hydroxylating *p*-coumaroyl-coA into 2’4’-dihydroxycinnamoyl-CoA. We offer tools for genomic and transcriptomic exploration in the context of jasmonate elicitation aiding in the characterization of novel stilbenoid-modifying and regulatory genes in the *Morus* genus.

## Introduction

Plants produce a myriad of specialized metabolites among which phenylpropanoids are one of the most widespread and abundant. These phenolic compounds have drawn increasing attention due to their multiple biological activities, preventing cardiovascular and neurodegenerative disorders or cancer and inflammatory-related processes, while also comprising strong antioxidant effects (Xu et al., 2017).

Within the phenylpropanoid pathway (PPP), stilbenes comprise a small group of phenolic compounds produced by the activity of stilbene synthases (STS) that evolved independently from chalcone synthases (CHS) in at least 72 unrelated plant species (Dubrovina & Kiselev, 2017; Rivière et al., 2012). Stilbenes accumulate in response to biotic stresses (Chong et al., 2009) and play important defense roles as phytoalexins against plant pathogens (Albert et al., 2011; Blanco-Ulate et al., 2015; Chalal et al., 2014), while they are also known to be produced upon ultraviolet radiation (Adrian et al., 2000; Kiselev et al., 2017; Z. Liu et al., 2019), and mechanical wounding (Chitarrini et al., 2017) via hormone signaling pathways mostly involving jasmonates (Billet et al., 2018). The synthesis of resveratrol by stilbene synthases (STSs) represents the first committed step of the pathway. Resveratrol can be stored inside the cell after being glycosylated into piceid (Hall & De Luca, 2007) but it is also transported out of the cells (Martínez-Márquez et al., 2023). Depending on the plant species, resveratrol can also be modified (i.e., methylated, acylated, hydroxylated and halogenated) or oligomerized (Valletta et al., 2021), with each derivative presenting different biological activities in humans (Navarro-Orcajada et al., 2022).

Plants from the Moraceae family have been largely used in traditional Chinese medicine due to their bioactive properties, mainly attributable to the presence of phenolic compounds including stilbenes. Species of the *Morus* genus, including mulberry trees (*Morus alba* L.), are known to accumulate several stilbenoids such as resveratrol, oxyresveratrol (oxyR, i.e., a hydroxylated form of resveratrol), and mulberroside A (mulA, an oxyR di-glucoside), in different organs of the plant, especially in hardwood and root bark (Zhou et al., 2013). With a considerable repertoire of pharmacological applications, *Morus alba* is now the most studied species for oxyresveratrol production (Lim et al., 2015). OxyR presents strong tyrosinase inhibitory activity (J.-K. Kim et al., 2012), together with anti-inflammatory (Thaweesest et al., 2022) and antioxidant (Heo et al., 2016) effects. Its protective properties against neurological diseases have also been reported (Ban et al. 2006).

The pathway for resveratrol biosynthesis in mulberry can be deduced from the current knowledge in grapevine (*Vitis vinifera* L.), a deeply studied high-stilbenoid producer with more than 40 *STS* genes in its genome (Vannozzi et al., 2018). A few *STS* genes have been isolated in Morus spp. (M. Li et al., 2016; Wang et al., 2017) but whether resveratrol is hydroxylated to form oxyR (as suggested by Zhou et al. (2013)) or if oxyR is produced in a different way (as suggested by S. Liu et al., 2022), this remains unsolved. Pathway genes responsible for the modification (e.g., monooxygenases, methyl- and glucosyl-transferases) or cellular export of resveratrol have not been yet identified in *Morus* spp. In addition to stilbenoids, other related compounds such as benzofuran derivatives (i.e., moracins) accumulate in mulberry tissues. Their biosynthetic pathway has not been demonstrated neither.

Despite the high value of stilbenoid compounds in mulberry, their natural production in plant tissues is limited and highly variable. Chemical synthesis or extraction from raw material, including non-renewable tissue, is technically complex, inefficient and unsustainable. Cell suspensions and root cultures arise as biotechnological solutions and can be combined with chemical or physical elicitation and metabolic engineering approaches. Stilbenes have been previously elicited and engineered in cell cultures of pine (Y. R. Kim et al., 2022), peanut (Vornam et al., 1988), pelargonium and grapevine berries (Belchí-Navarro et al., 2012; Lijavetzky et al., 2008; Martinez-Esteso et al., 2011; Martínez-Márquez et al., 2016). Inyai et al. (2021) achieved high amounts of oxyR and mulA in *M. alba* hairy root cultures (HRs) by using methyl jasmonate and yeast extract, while Fang et al. (2022) combined methyl jasmonates, methyl-β-cyclodextrins and magnesium chloride as elicitors in HRs. Till now, this combined elicitation has not yet been accomplished using fully-suspended cell cultures of mulberry. Instead, it has only been performed using plantlets (Sabater-Jara et al., 2023) or in hydroxypropyl beta cyclodextrin-treated calli (Komaikul et al., 2019).

In addition to plant cell and tissue culture, metabolic engineering of microorganisms represents a powerful means for industry-scaled production of oxyR, overcoming the limitations of plant-based production. This approach is gaining much interest to produce stilbenes (Jeandet et al., 2018; Shrestha et al., 2019). In this context, the identification and functional characterization of the genes responsible for oxyR production becomes crucial. A first hypothesis is that oxyR synthesis involves hydroxylases some of which correspond to cytochrome P450 (CYP450) monooxygenases. Whether these act before or subsequently after resveratrol is produced, is unknown. Hydroxylases have undergone extensive duplication events throughout the evolution of plant genomes (e.g., there are 174 CYP450 genes identified in *Morus notabilis* (Ma et al., 2014). Selecting genes for their later characterization through biochemical or biotechnological methods becomes extremely laborious and demands for genome-wide approaches to narrow down the list of potentially related genes. Here, we initiated mulberry cell suspensions to perform elicitation experiments and generate time-course multi-omics datasets to identify and further characterize oxyresveratrol-producing enzymes. Our data pinpoints to the involvement of p-coumaroyl CoA 2’-hydroxylase enzymes acting upstream of stilbene synthases.

## Results

### Methyl Jasmonate and cyclodextrins elicit oxyresveratrol and resveratrol production in mulberry cell suspensions

Oxyresveratrol and piceatannol correspond to the meta- and ortho-hydroxylated derivatives of resveratrol, respectively. This high similarity makes their chromatographic separation difficult. In order to determine whether mulberry was able to produce both positional isomers, we first explored analytical methods that could discriminate between them. Following the MRM methodology development of Hurtado-Gaitán et al. (2017) we were able to find a qualifier fragment for oxyresveratrol, of [M+H^+^]=161 (Figure 1A; Supplementary Figure 1A), which allowed differentiating both isomeric compounds.

Cell suspensions of *Morus alba* (Figure 1B) were initiated from undifferentiated calli established from twigs of specimens belonging to white- and pigmented-fruit cultivars (Supplementary Figure 1B). These were treated with methyl jasmonate (MeJA) and methyl-β-(MBCD) or 2-Hydroxypropyl-β-(HPBCD) cyclodextrins. Stilbenes emit violet-blue fluorescence under UV light radiation (Poutaraud et al., 2007). As expected, elicited mulberry cell cultures showed this fluorescence when exposed to ultraviolet light (312 nm; Figure 1C). We observed an increase in the accumulation of resveratrol and oxyresveratrol at 4-5 days post-elicitation (Supplementary Figure 1C-D, Supplementary Tables 1-2), while piceatannol was not detected. The treatment with methyl jasmonate alone had a negligible effect on the production of these compounds, while the treatment with MBCD alone showed a moderate effect. Compared to the single treatments, the combined effect of MBCD and MeJA led to a substantial increase in the synthesis of both resveratrol and oxyresveratrol. Suspensions derived from the pigmented cultivar produced slightly less resveratrol when compared to white cultivar-derived cells. However, a greater accumulation of oxyresveratrol was observed (Supplementary Figure 1D). Therefore, cell suspensions generated from the pigmented fruit cultivar were chosen for further analysis.

A time-course experiment was performed with a combined treatment of elicitors (MBCD-MeJA) with sampling following at different time points: 0.5, 6, 12, 24, 48 and 72 hours. Both the intra- and extracellular production of oxyresveratrol, which was considerably higher than that of resveratrol, increased steadily until reaching its maximum at 48 hours, where it remained constant (Figure 1C; Supplementary Table 3). Extracellular resveratrol, instead, increased at a constraint ratio and didn’t seem to reach a plateau at any of these time points. The quantification of stilbene glycosides revealed that mulberroside A experienced a reduction in response to elicitation, especially in the latest time-points. The accumulation of piceid was almost insignificant, despite a slightly higher accumulation in elicited cells in almost all time points.

**Figure 1.**
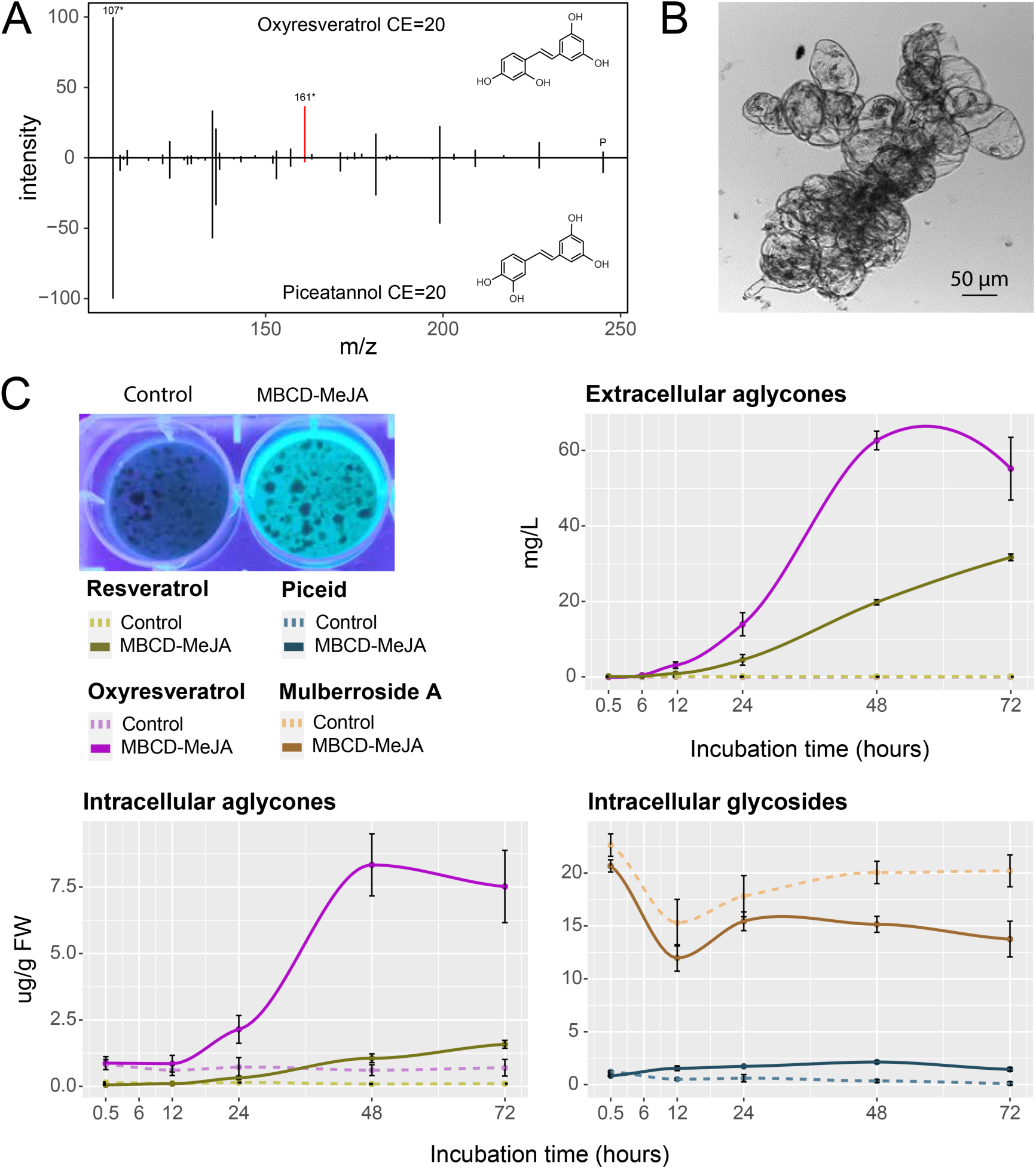
The combined treatment of MeJA and MBCD promotes the accumulation of resveratrol and oxyresveratrol in mulberry cell suspensions. A) Mass-spectrometry distinction of mono-hydroxylated derivatives of resveratrol (i.e, piceatannol and oxyresveratrol). Mirror plot showing fragmentation mass spectra of oxyresveratrol (top) and piceatannol (bottom) at collision energy (CE) of 20eV. m/z 107 and m/z 161 correspond to the quantifier and qualifier ion fragments, respectively. P stands for precursor ion at m/z 245 B) Bright field image of untreated cells grown in liquid media. C) Blue fluorescence emitted after UV-C irradiation of control and elicited cell suspensions (top-left panel), and time-series accumulation of extra and intracellular stilbene aglycones (and intracellular glycosides) in response to methyl jasmonate and cyclodextrin elicitation. Intracellular concentrations are given in ug/g of fresh weight (FW) while extracellular accumulation is measured in mg/L. Data represent mean ± sd.

### Reannotating the mulberry genome for the identification of stilbenoid-related genes

Despite the well-documented accumulation of resveratrol and oxyresveratrol in mulberry, the current annotation of its genome (20,386 genes; Jiao et al. (2020)) fails to harbor any stilbene synthase (*STS*) gene. This annotation seemed therefore incomplete, supporting the need for a more accurate annotation to allow further functional characterizations of the genes involved in elicited pathways. We followed a reannotation workflow to define a more complete set of protein-coding genes using an automated pipeline that integrated protein homology, short and long-read transcript evidence and *ab initio* predictions. A total of 85,569 transcripts were obtained when combining the official and the updated annotations. Filtering out redundancy resulted in 31,401 gene models. Gene space completeness was overall improved, with BUSCO results showing that 95.7% of the universal single-copy genes from the eudicots clade were present, compared to 92.1% from the official annotation (Figure 2A). While the total number of genes considerably increased, the number of single-copy genes in more than one copy were also decreased (from 24.78% to 5.35%).

The structural annotation was followed by a gene function prediction pipeline based on the MapMan ontology. This allowed for the identification of gene families belonging to the shikimate and early phenylpropanoid pathways, somewhat comparable in copy number to those in grapevine (Figure 2B). We observed a few *STS* genes models wrongly annotated as chalcone synthases while the rest were not even present in the official annotation (Supplementary Figure 2). We observed *STS* genes located on chromosome 8 in two distinct genomic clusters (Figure 2C). The evidence of such a large duplication event was unique in the entire chromosome. In addition, the two regions comprise a similar number of *STS* genes suggesting that one of them was caused by a segmental duplication event of the other cluster (Supplementary Figure 3). A total of 22 *STS* genes were identified in this chromosome, with 12 in the first cluster and 10 in the second. Half of the genes seem to correspond to pseudogenes, despite this is only based on the presence of premature stop codons or absence of start/stop codons, and not in relation to expression behavior.

Several genes were manually curated using the Apollo interface, guided by both the short and long-read sequencing data generated in this study. To this purpose, we also included public data obtained from fruit and leaf samples (Illumina-sequenced libraries). The phylogenetic reconstruction of the polyketide synthase family, using these manually curated gene models together with grapevine sequences, confirmed that several genes that were previously annotated as *CHS* clearly clustered separately from *V. vinifera CHSs*. STS, CHS and PKSIIIA/PKSIIIB proteins clustered distinctly depending on each group, with all CHS sequences falling under a single group with all species, while STSs clustered separately for each species (Figure 2D). This observation is consistent with a convergent evolution for *STS* genes as suggested by Tropf et al. (1994). Finally, PKSIII-like proteins seem not present in *M. alba*.

**Figure 2.**
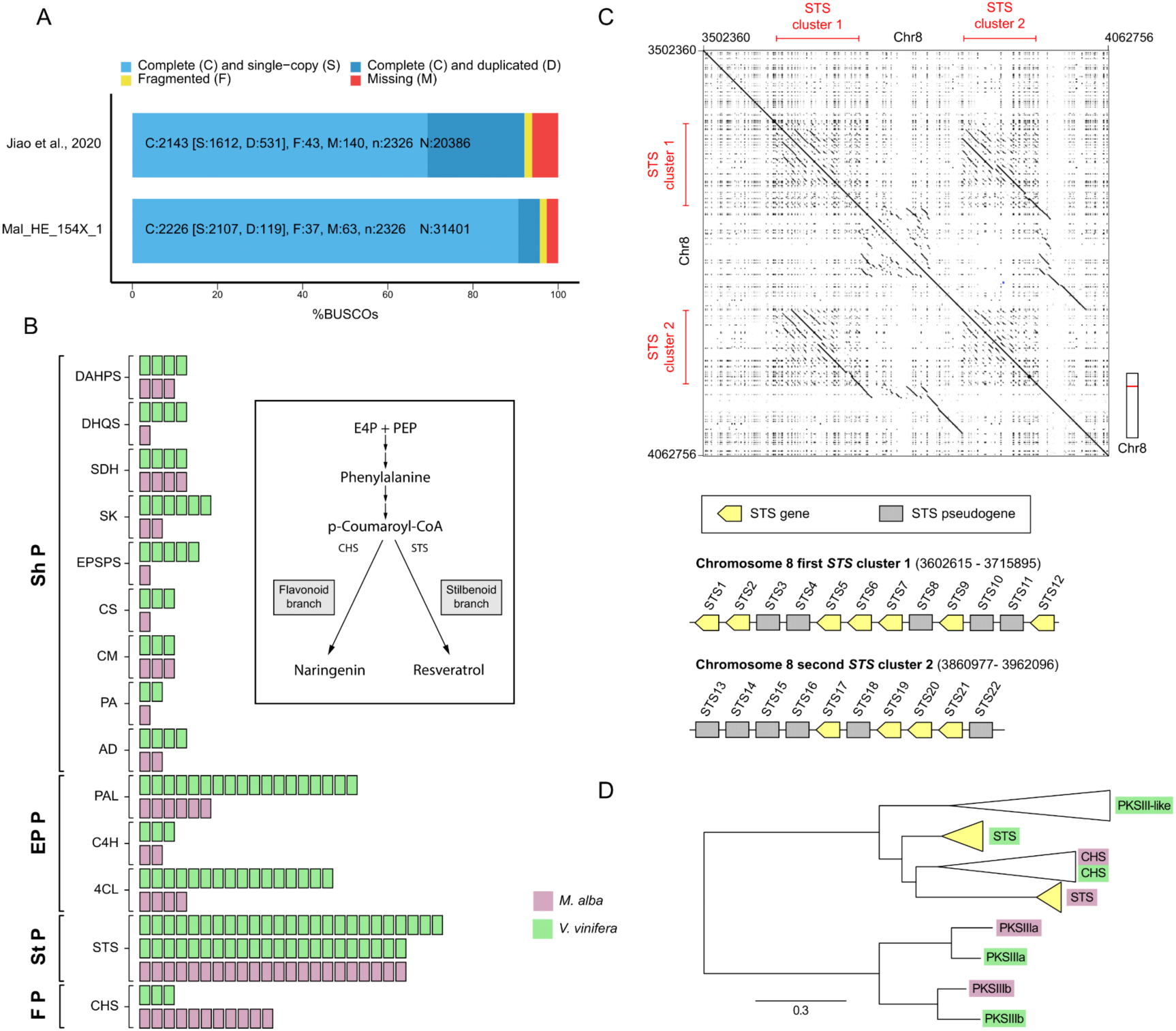
An improved annotation of the *M. alba* genome reveals two *STS* gene clusters located in chromosome 8. A) BUSCO completeness analysis for comparing the official (Jiao et al., 2020) and updated structural annotations based on the eudicots dataset. Colors represent the different classes of the BUSCO assessment results. n= total number of gene models in BUSCO dataset, N= total number of gene models in annotation. B) Comparison between the number of enzymes belonging to the shikimate (ShP), early phenylpropanoid (EPP), stilbenoid (StP) pathways in *V. vinifera* and *M. alba*. Chalcone synthases (CHS) from the flavonoid pathway (FP) are also included). C) Dotplot of chromosome 8 and microsynteny analysis of *STS* clusters showing segmental duplications forming paralogous sets of genes (top panel). Genomic organization of *STS* genes annotated in the *M. alba* reference genome (bottom panel). Some genes are classified as pseudogenes due to the presence of premature stop codons in their coding sequences (CDS). D) Maximum likelihood phylogenetic tree of the polyketide synthase III (PKSIII) family, containing STS, CHS, PKSIII-like, and PKSIIIa/b proteins from *V. vinifera* and *M. alba*. The tree corroborates independent evolutionary origins of stilbenoid synthases in both species. Evolutionary distances are represented as amino acid substitutions per site. The tree was constructed using 1000 bootstrap values (values found in the uncondensed tree, Supplementary Figure 4).

### Multi-omics integration to identify and further validate *C2’H* involved in oxyR synthesis

The gene expression profiles of the *M. alba* cell cultures treated with MBCD-MeJA were analyzed using weighted gene co-expression network analysis (WGCNA) providing a method to cluster genes into modules based on their expression similarity. In addition, we also correlated these modules with metabolite abundances. WGCNA resulted in a free-scale topology network with a total of 44 modules, with most *STSs* found in module 39, followed by modules ME7 and ME11 (Figure 3A).

**Figure 3.**
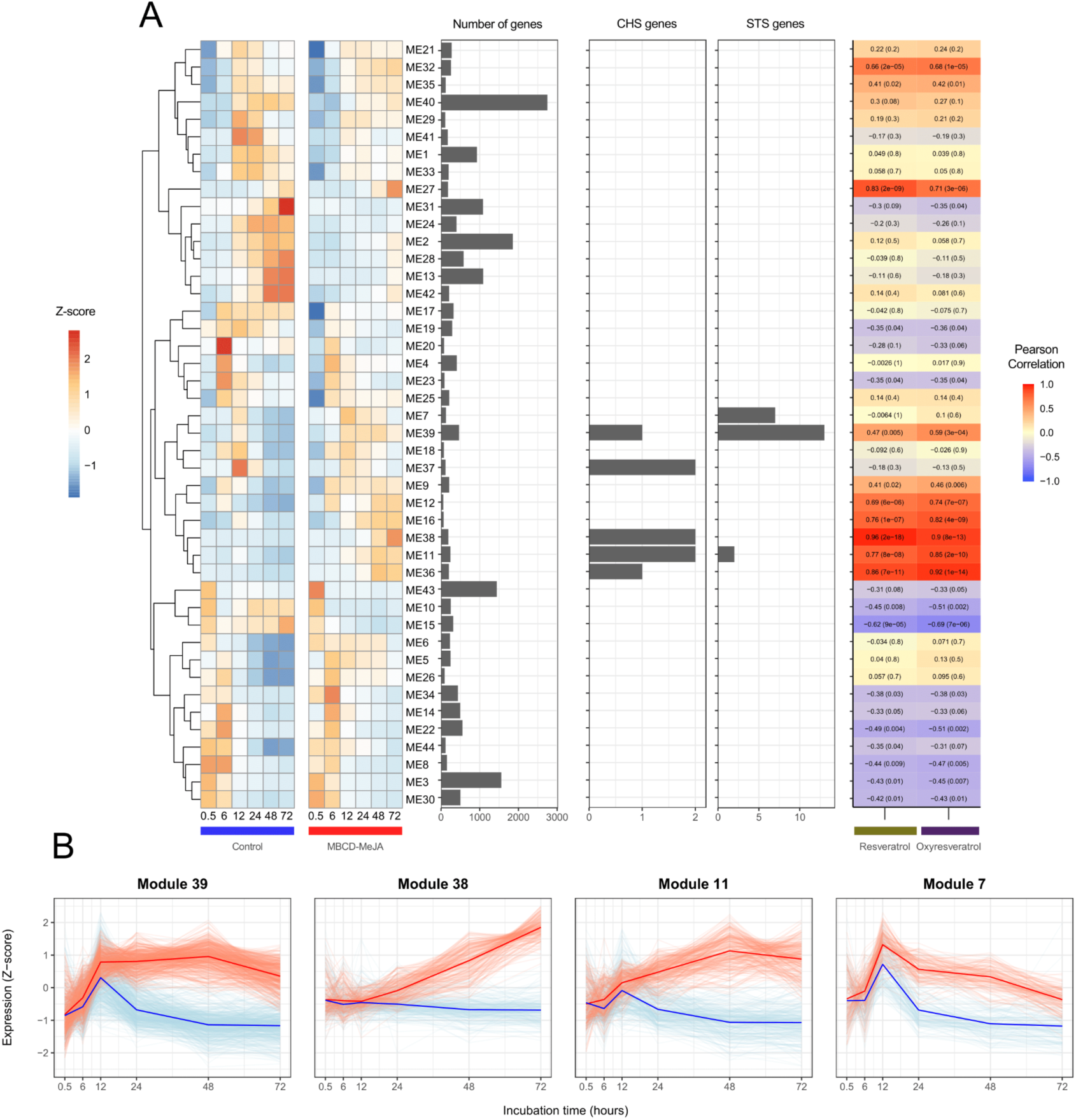
Co-expression analysis and metabolomic/transcriptomic integration to identify potential oxyresveratrol biosynthetic genes. A) Time-related clusterization of the elicited transcriptome of *M. alba* by Weighted Gene Co-expression Network Analysis (WGCNA) (extreme left). Correlation of each module eigengene (ME) with the time-course accumulation of resveratrol and oxyresveratrol (extreme right). Values inside each colored box represent the correlation coefficients and *p*-values (in parenthesis) obtained between each module and the respective metabolite. Bar plots depict the total number *CHS* and *STS* genes per module. B) Individual gene expression profiles for a selection of modules showing activated transcript profiles in response to elicitation. FPKM data is shown scaled (Z-score). Dark color trajectories on each module represent the two mean gene expression profiles for each condition. Red and blue lines correspond to elicited and control samples, respectively.

The selected modules shared a similar expression pattern with up-regulation in response to elicitation (Figure 3B). Interestingly, some *CHSs* were clustered in these modules too. Module ME11, with several *STS* and *CHS* genes inside, showed a high and significant correlation with the accumulation of resveratrol and oxyresveratrol. Module ME39 showed less correlation, but still significant. Module ME38, despite not possessing any *STS* gene, presented the highest correlation with both resveratrol and oxyresveratrol.

The functional enrichment analysis of these four modules revealed a few additional significant MapMan categories with genes different from those conferring CHS/STS activity (Supplementary Table 4). This evidence suggested their participation in the stilbenoid pathway following the *guilty-by-association* principle. Among these, we found the term feruloyl-CoA 6-hydroxylase (F6’H) as overrepresented in modules 39 and 7, where *STSs* were also present (Figure 4A). As in the case of *MalSTSs*, a few of the mulberry genes from this group had also been revealed following the genome re-annotation; Supplementary Figure 5). On the other hand, the cytochrome P450 hydroxylase term was enriched in modules 38 and 11.

To further investigate the potential F6’H gene family being highly co-expressed with *STS* genes, we conducted a phylogenetic analysis that included F6’Hs from *A. thaliana* and *V. vinifera*, in addition to a group of enzymes (named Ib2) from sweet potato (*Ipomoea batatas*) that derive from *F6’H* genes and catalyze the ortho-hydroxylation of *p*-coumaroyl-CoA (C2’H activity; (Matsumoto et al., 2012). The inclusion of C3’H and C4’H proteins from *A. thaliana*, *V. vinifera* and their homologues found in *M. alba*, revealed two distinct groups (Supplementary Figure 6). A first group clustered F6’H proteins from Arabidopsis and grapevine and only one from mulberry, while the other clade contained the remaining six mulberry F6’H sequences being co-expressed with STSs.

Umbelliferone (7-hydroxy-coumarin) is one of the end-products of C2’H activity in sweet potato (Matsumoto et al., 2012), produced by the reduction of hydroxylated p-coumaroyl-CoA. On the contrary, F6’Hs are only able to produce scopoletin after the hydroxylation of feruloyl-CoA. If any of the elicited *MalF6’H*-like genes encodes for an intrinsic C2’H enzymatic activity, umbelliferone should be found in elicited cells. We in fact observed its unique accumulation upon elicitation as evidenced through GC-MS analysis of the extracellular medium (Supplementary Figure 7). Mulberry sequences within the separate F6’H clade were thus hypothesized to be acting as C2’Hs rather than actual F6’Hs, despite still retaining 2-oxoglutarate (2OG) and Fe(II)-dependent oxygenase superfamily domains. The particular differences in the MalC2’H protein sequences could be responsible for a substrate switch from feruloyl-CoA to p-Coumaroyl-CoA as reported in sweet potato by Sun et al. (2015). Moreover, we hypothesized that the production of hydroxylated p-coumaroyl-CoA in elicited conditions could be a substrate of STSs for catalyzing the production of oxyresveratrol.

The expression of *STS* and putative-*C2’H* genes showed a peak at 12 hours post elicitation and also correlated with a general activation of the shikimate and early phenylpropanoid pathways (Supplementary Figure 8). In line with the subtle correlation between the accumulation of oxyresveratrol, resveratrol and *STS*-containing modules, we observed a delayed metabolite accumulation compared to the gene expression profiles. Considering the full trajectory of gene expression and metabolite accumulation, *STS* and *C2’H* genes initiate a decrease in their expression at 48h despite stilbenes still accumulating (Figure 4B). We further investigated the relationship between protein abundance and mRNA expression in the elicited cells, in order to predict whether despite their decrease in expression, *STS* and *C2’H* mRNAs were still being translated after 48h. We analyzed the three latest time points from control and elicitation samples using a proteomic label-free quantification approach to capture the elicited cell proteome changes. We identified a total of 1891 protein groups corresponding to 3158 unique proteins. From these, 712 proteins were considered as differentially abundant (DAP) in at least one elicitation time-point when compared to the control. For a total of 2793 mRNA-protein matching pairs, more than 60% showed either a positive or a negative correlation between their transcript/protein levels. 823 mRNA-protein pairs showed a positive correlation (Spearman’s rank coefficient ρ ≥ 0.4), with 134 being significant (ρ ≥ 0.8 and p-value < 0.05), while 910 showed a negative correlation (ρ ≤ -0.4), with 106 being significant (ρ ≤ -0.8 and p-value < 0.05). In addition, 1060 pairs did not show a positive nor a negative correlation between their mRNA and protein abundances. Each mRNA-protein pair was classified according to their differential expressions and abundances, resulting in 246 DEG (Differentially expressed genes)-DAPs (Differentially abundant proteins), 522 DEG-NVPs (Non variant proteins), 376 NVG (Non variant genes)-DAPs, 1649 NVG-NVPs. The median ρ for the complete data set (n = 2793) was -0.08, but this value varied for each one of the DE/DA subsets. DEG-DAP pairs had the highest ρ (median ρ = 0.7), indicating a stronger positive correlation between protein abundance and RNA expression for these pairs. In the case of DEG-NVP and NVG-DAP pairs, there was a higher proportion of negatively correlated mRNA-protein pairs than in the other subgroups (Figure 4C). A second WGCNA analysis of the mRNA-protein pairs showed that the most abundant modules corresponded to those DEG-DAP pairs induced upon elicitation (Figure 4D). Among these, STS and C2’H proteins showed very high correlation with their corresponding mRNAs in all cases (Figure 4E). This was certainly evidenced when comparing control and elicited conditions. However, while mRNAs of *STS* and *C2’H* tend to decrease towards the latest time point, protein abundances are kept constant or show a slight increase. This also suggested a higher correlation between these proteins and the abundances of oxyresveratrol and resveratrol than their mRNA abundances.

To validate whether the putative C2’Hs could be involved in oxyresveratrol biosynthesis we transiently expressed *C2’H1*, *C2’H4* and *C2’H5,* in combination with a previously characterized *STS* (Santos-Rosa et al., 2008), in *Nicotiana benthamiana* leaves. The single and combined effects of *C2’H* and *STS* agroinfiltration were assessed by LC-MS at 3- and 6-days post agroinfiltration for the quantification of stilbenoids. Several peaks were detected in the corresponding ion transition channels of oxyresveratrol and resveratrol but with similar retention times as their glucosides (i.e., mulberroside A and piceid, respectively). This strongly suggests that non-specific mono-glycosylated forms of oxyresveratrol are being produced in all *STS*+*C2’H* combinations, in the same way as the pure infiltration of oxyresveratrol in leaves (Figure 4F), all of which suffer partial *in-source* fragmentation. In a comparable fashion, the infiltration of the single *STS* or *STS*+*C2’H* constructs generated peaks in the resveratrol channel matching the retention time of piceid. Neither the infiltration of C2’H alone or C2’H plus resveratrol produced any of the potential oxyresveratrol-glycosylated forms, suggesting that C2’Hs require STS action but do not hydroxylate resveratrol to form oxyresveratrol.

To directly demonstrate the production of the oxyresveratrol aglycone by the subsequent activity of C2’Hs and STSs, we transiently expressed these genes in cell suspensions of *Vitis vinifera* cv. ‘Gamay’ (Figure 4G), which generally accumulate aglycones under elicitation (Orduña et al., 2022). Oxyresveratrol was detected within intracellular extracts of cells transiently transformed with *STS* and *C2’H*, at 6 days after transient transformation. Altogether, our results demonstrate the requirement of *p*-coumaroyl-CoA 2’-hydroxylases in the synthesis of oxyresveratrol in *Morus alba*.

**Figure 4.**
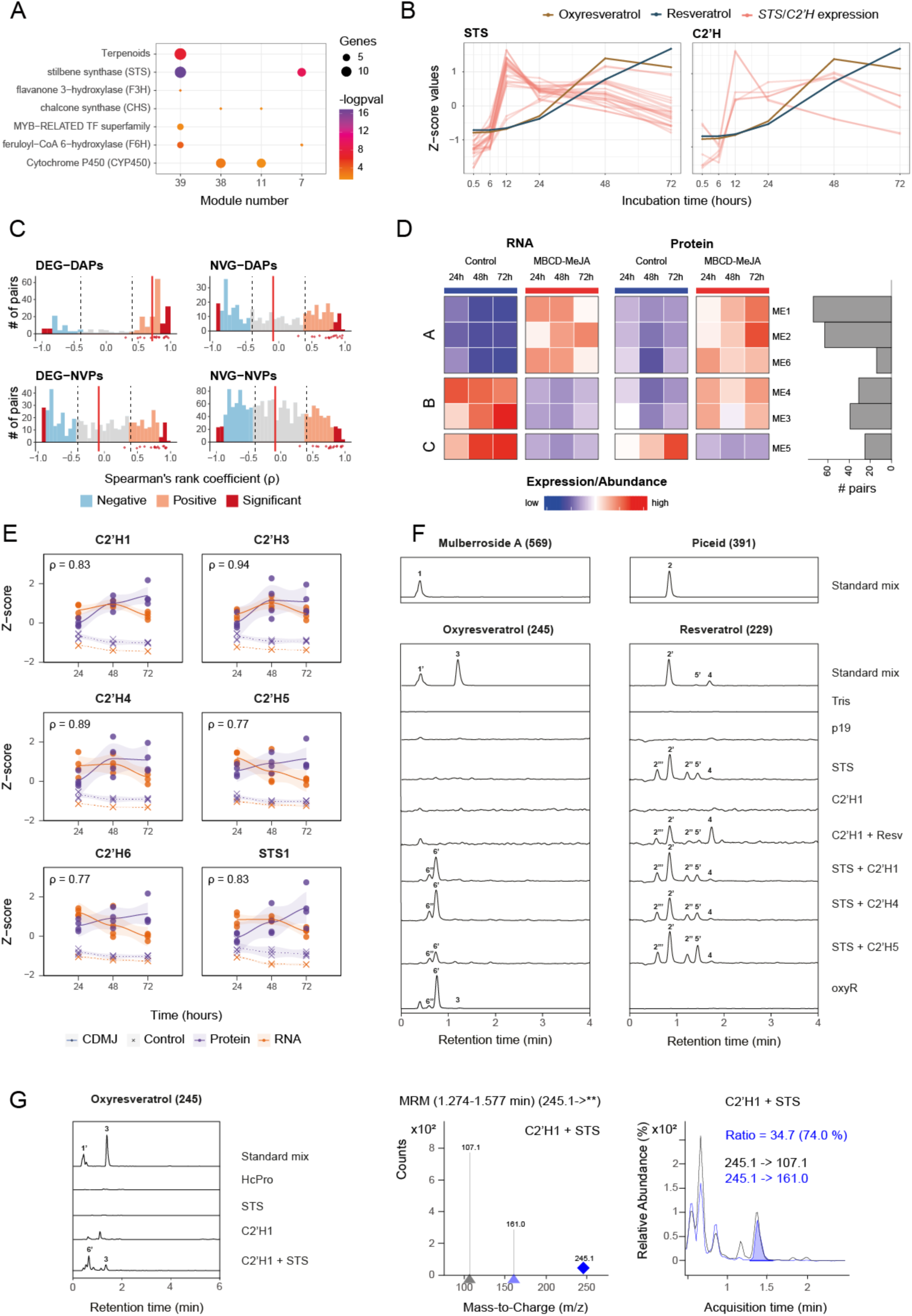
C2’H enzymes are involved in the production of oxyresveratrol and respond to elicitation. A) Selection of significantly enriched MapMan terms in selected WGCNA modules. The number of genes intersecting with each ontology term is represented by dot size. B) *STS* and *C2’H* gene expression profiles compared to resveratrol and oxyresveratrol abundances. C) Spearman’s rank coefficient (ρ) values of mRNA-protein pairs according to their stage differential expressions (DE)/abundances (DA) DEG-DAPs, DEG-NVPs, NVG-DAPs and NVG-NVPs (NV: non-variant). Red lines represent the median correlation for each subset. Dashed lines indicate the limits to consider positive and negative correlations. Red dots under the *x* axis refer to proteins belonging to the phenylpropanoid pathway identified in Figure 2B. D) Z-scored abundances of stage variant mRNA/protein pairs clustered by WGCNA. E) Abundances and correlations of C2’H mRNA/protein pairs putatively involved in oxyresveratrol synthesis (other related pairs can be found in Supplementary Figure 9). F) LC-Mass Spectrometry chromatograms obtained from methanol extracts of agroinfiltrated *Nicotiana benthamiana* leaves. Leaves were co-infiltrated with *Agrobacterium tumefaciens* harboring the P19, *STS* or *C2’H* genes, in different combinations. Resveratrol and oxyresveratrol infiltrations are also included together with Tris buffer, the latter also used as a negative control. Ion current data was extracted for each ion transition (channels) of resveratrol, oxyresveratrol and their glycosylated forms in the MRM positive mode. Standard mix includes resveratrol, oxyR, piceid and mulberroside A. Each chromatogram is representative of three to six independent infiltrations carried out in different plants and which can be observed in Supplementary Figure 10. Selected peaks: 1-Mulberroside A, 2-trans-piceid, 2’-trans-piceid (SF), 2’’, 2’’’-resveratrol glycosides (SF), 3-oxyresveratrol, 4-resveratrol, 5’-cis-piceid (SF), 6’,6’’-oxyresveratrol glycosides (SF). SF: in-source fragmentation. G) MRM analysis of oxyresveratrol within intracellular extracts of grapevine cell suspensions transiently transformed with *HcPro*, *STS* or/and *C2’H*, at six days after transfection. Left: Extracted ion chromatograms of transition from oxyresveratrol (245>107; more detail in Supplementary Figure 11). Center: composite MS/MS spectrum of the recorded transitions, indicating the precursor ion m/z with a diamond. Right: normalized chromatographic traces of the monitored transitions for oxyresveratrol.

## Discussion

Cell suspension cultures are a promising way to obtain valuable plant natural compounds. This becomes particularly relevant for molecules whose utility in society is affected by low efficient and unsustainable plant extraction methods or in the use of chemical synthesis that produces non-specific mixtures and generates toxic waste with high production costs.

In this study, we show that mulberry cell cultures produce large amounts of both resveratrol and oxyresveratrol when elicited with methyl jasmonate and cyclodextrins. The amounts generated seem appealing for potential applications in pharma and cosmetic industries. These production rates resemble those of hairy roots been elicited with the same type of compounds (Fang et al., 2022). Nonetheless, as shown while comparing pigmented- and white-fruited cultivars, differences in stilbenoid production can be found depending on the starting material used for the establishment of the cell suspension. A synergistic action of jasmonates and cyclodextrins have already proven to trigger an enhancement of stilbenoid production in grape cell suspensions (Almagro et al., 2014; Halder et al., 2019). The constitutive secretion of these compounds into the growing media could be exerted through passive or active transport, via transporters such as those identified in grape (Martínez-Márquez et al., 2023) or via the intrinsic capacity of cyclodextrins acting as carriers (Morales et al., 1998). This secretion facilitates their extraction and purification without affecting the cells in the suspension.

We demonstrated that elicitation boosts the expression of enzyme-coding genes involved in primary (i.e., shikimate) and specialized (early phenylpropanoid and stilbenoid) metabolic pathways. However, the lack of genes belonging to the stilbenoid pathway in the official *M. alba* genome annotation (Jiao et al., 2020) prompted us to integrate transcriptomic resources to revise this resource, especially in order to ascertain the presence of *STS* genes. The *de novo* transcriptomes, assembled with long-read sequencing data, also allowed us to ensure that the new gene models were correctly annotated. Our gene set was more complete specially in the biological context of specialized metabolism and its responses to methyl jasmonate.

Two distinct *STS* clusters were found in chromosome 8, probably originating from different duplication events. The observed sequence similarity patterns suggest that a first cluster of *STS* genes was initially formed through tandem duplication events, which were later followed by a segmental duplication of the first cluster. Such an expansion could be the result of a positive selection pressure, indicating the importance of stilbene accumulation in the fitness and/or survival of *M. alba*. Resveratrol and its derivatives (e.g., its methylated, glycosylated, acylated forms) are in fact known as phytoalexins, capable of protecting the plant against reactive oxygen species and pathogens, e.g. *Botrytis cinerea* (Adrian et al., 1997). In addition, oxyresveratrol has been found to be a powerful free radical scavenger, surpassing the capacity of resveratrol (Lorenz et al., 2003).

Chalcone and stilbene synthases compete for p-coumaroyl-CoA and malonyl-CoA substrates. This is due to the fact that both enzymes are polyketide synthase type III and in fact STS activity evolves from a CHS ancestor. Our phylogenetic analysis was able to group all CHS sequences together. However, STSs from grapevine and mulberry clustered independently, an expected outcome due to the known convergent evolution of STSs. STS have emerged in more than 70 unrelated plant species. Future studies can focus on the characterization of the entire STS family in mulberry, to prove their activity, but also to understand the conformational effects of different mutations in CHS leading to STS activity in plants. Our transcriptomic analysis showed that both *STS* and *CHS* genes were induced upon jasmonate-mediated elicitation. This suggests that both stilbene and flavonoid pathways are induced. Other closely related pathways such as the benzofuran family of compounds (e.g., moracin) are also induced (Fang et al., 2022). Our data and resources can guide gene discovery approaches to identify the genes responsible for these other pathways.

It has normally been accepted that resveratrol is the first committed compound of the stilbenoid pathway. Under this paradigm, resveratrol derivatives normally occur subsequently as resveratrol-modifying enzymes come into play. This is the proven case in resveratrol glycosylation, prenylation, acylation and methylation (reviewed by Valletta et al. (2021)) but it has also been the hypothesis for the hydroxylation of resveratrol into piceatannol (Martínez-Márquez et al., 2016). In the case of oxyresveratrol, it has been hypothesized to be originated from the oxidation of resveratrol (Zhou et al., 2013), till now, where we prove wrong.

Coumarins are formed by the 2/6-hydroxylation of the *p*-coumaroyl phenyl ring followed by trans/cis isomerization of the side chain and lactonization. The first step of the synthesis of the 2H-1-benzopyran-2-one structural core of coumarins of has been demonstrated to be enzymatically catalyzed, whilst the subsequent steps may occur spontaneously under light conditions or enzyme catalyzed (Stringlis et al., 2019; Vanholme et al., 2019). In *A. thaliana*, feruloyl CoA 6’-hydroxylase (*At*F6’H) catalyzes the committed step for the synthesis of the coumarin scopoletin, displaying high specificity (Kai et al., 2008). A *p*-coumaroyl CoA 2’-hydroxylase of *Ruta graveolens* (*Rg*C2’H) catalyzes the committed step for the synthesis of umbelliferone (Vialart et al., 2012). Interestingly, RgC2’H is also able to hydroxylate the equivalent position of feruloyl CoA (F6’H) to promote the synthesis of scopoletin. The induction of C2’H in elicited cells of *M. alba* is thus consistent with the production of umbelliferone in response to methyl jasmonate and cyclodextrins. Scopoletin however was not detected, suggesting either that feruloyl CoA is absent in *M. alba* cells or that *Ma*C2’H displays a high specificity for p-coumaroyl-CoA. Unexpectedly, the use of the C2’H product 2’,4’-dihydroxycinnamyl CoA by STS provides a plausible pathway for the production of oxyresveratrol in *M. alba*.

Our findings suggest that C2’H enzymes synthesize 2’4’-dihydroxycinnamoyl-CoA from p-coumaroyl-CoA which is in turn the substrate for STS to convert oxyresveratrol (Figure 5). Under this model, STS enzymes can accept both hydroxylation states of p-coumaroyl as a substrate. We also suggest that the enzymatic activity of C2’Hs in terms of velocity and performance must be high, as oxyresveratrol production rates are higher to those of resveratrol under elicitation. The spontaneous production of umbelliferone is also probably slow and has a minor effect of oxyresveratrol levels.

**Figure 5.**
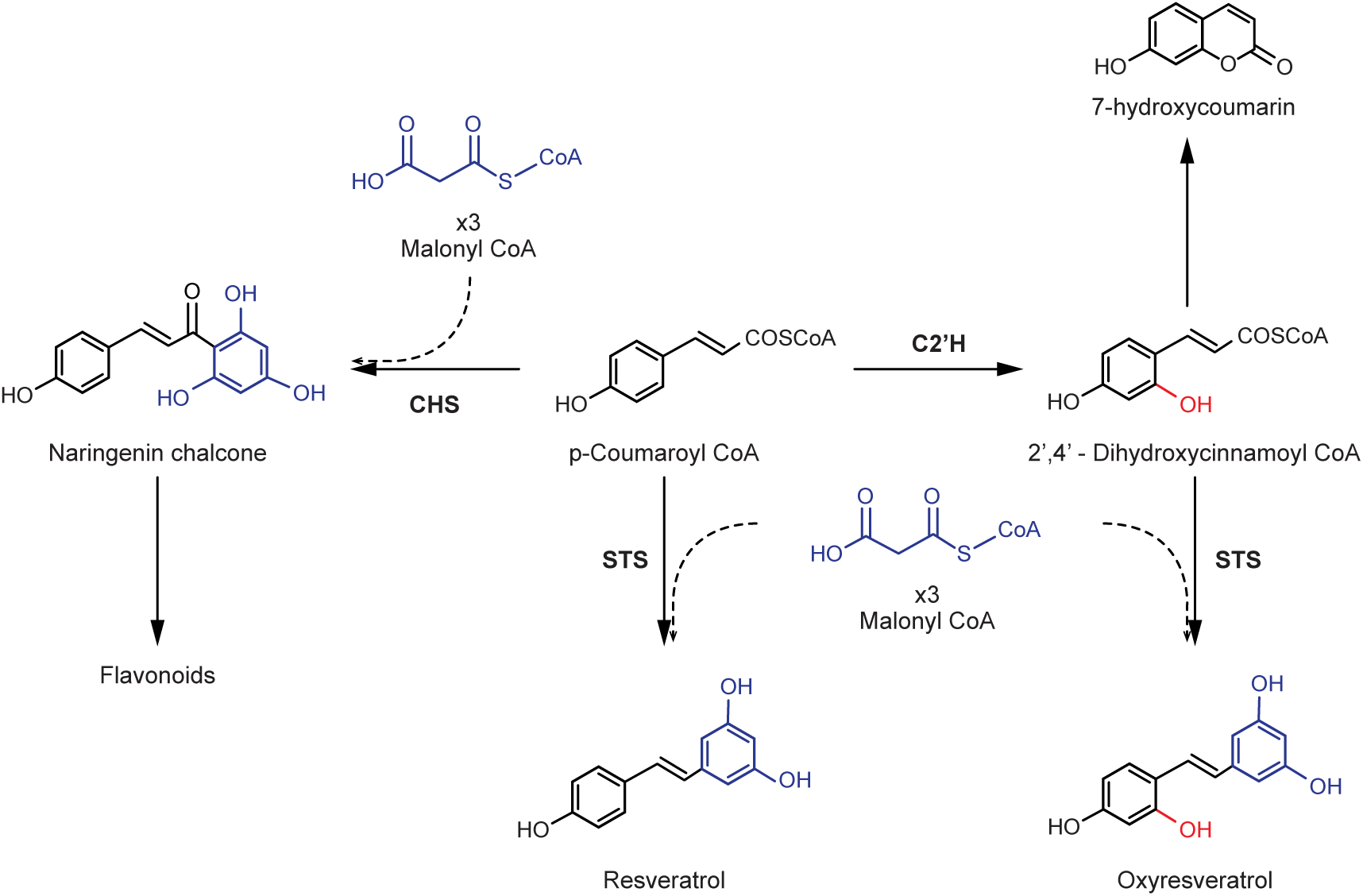
Oxyresveratrol biosynthetic pathway as revealed by integrative multi-omics and functional analyses. Resveratrol and oxyresveratrol are both produced by STS activity acting in parallel on p-coumaroyl-CoA and di-hydroxycinnamoyl-CoA as substrates, respectively. The proposed model refutes previous hypotheses that all stilbenoid forms are derived from a resveratrol backbone. STS: stilbene synthase, CHS: chalcone synthase; C2’H: p-coumaroyl CoA 2’-hydroxylase. Umbelliferone (7-hydroxycoumarin) is also produced from di-hydroxycinnamoyl-CoA.

We provide experimental evidences using heterologous plant systems that *C2’H* expression is required to produce oxyresveratrol. The well-known fact that the stilbene core can be formed from substrates different from p-coumaroyl CoA, such as cinnamoyl CoA or caffeoyl CoA (Raiber et al., 1995) and the occurrence of STS as gene families (Vannozzi et al. (2012), Figure 2D) as potential source of differential specificity, provide arguments that support a biosynthetic pathway for oxyresveratrol parallel to that of resveratrol instead of a linear pathway of resveratrol hydroxylation in *M. alba*.

We cannot imply that the production of other hydroxylated stilbenes, such as piceatannol, is led in the same way as for oxyresveratrol. In fact, grapevine doesńt seem to have C2’H activity and Martínez-Márquez et al. (2016) showed that a human cytochrome P450 enzyme was capable of producing piceatannol when agroinfiltrated together with *STS* genes in *N. benthamiana*. However, our work contributes to a new paradigm where STS enzymes can be promiscuous for different types of substrates. In line with this, STSs from pine (Pinus sp.) can use cinnamoyl-CoA to produce pinosylvin (Schanz et al., 1992).

The expression and abundance patterns of STS and C2’H genes and proteins correlate well with the accumulation of resveratrol and oxyresveratrol. This suggests that these genes are co-regulated and act in a coordinated manner to produce these compounds. The tight co-expression of these genes suggest that they may be controlled by the same transcriptional regulators. In grapevine, Subgroup 2 R2R3-MYBs are able to control the expression of almost the entire *STS* gene family, while also directly regulating shikimate and early phenylpropanoid pathway genes.

Our open access resources act as repositories of novel genes to study in their relation to stilbene production in mulberry. We have centralized all the data generated in this work. Specifically, the new gene annotation or transcriptomic data can be downloaded or explored from MorusalbaDB and the Mulberry Visualization Platform (MulViz), respectively. These are the first open access databases created for this species. In this, resource users will be able to browse the genome, explore the functional and structural information of the annotated genes, as well as browse gene models in the genome and visualize co-expression data. Despite these advances, some gene families may still require manual curation, for which we have also created a curation interface via Apollo where the community can access and contribute for the best annotation of mulberry.

## Methods

### Initiation, growth conditions and elicitation of Morus cell suspensions

*Morus alba* undifferentiated cell growth in the form of calli was achieved using young stems (twigs) cut into 2 cm length pieces as explants by Mr. Juan J. Montesinos. These were collected in 2012 from two specimens located at the campus of University of Alicante (Spain), one with intensely pigmented and another with non-pigmented fruits out of which we obtained a line named “red” and another named “white”, respectively (Supplementary Figure 1B). Both were assessed to be *M. alba* on the basis of both ITS length and ploidy according to Zeng et al. (2015). After thorough surface sterilization (sequential rinses in sterile water 3 times, once in 70% ethanol for 30 sec, water 3 times, once in 1% sodium hypochlorite for 3 min and final rinses in sterile water) explants were wiped with sterile filter paper, cut longitudinally into two halves and placed with the cutting surface onto solid medium with different hormonal balance. Medium composition per liter was 4,3 g Murasighe and Skoog basal medium, 30 g sucrose, 0.25 g casein hydrolysate, 1 mg calcium pantotenate, 100 mg myoinositol, 0.01 mg biotin, 1 mg pyridoxin, 1 mg thyamine, 1 mg nicotinic acid, 7.5 mg FeSO_4_.7H_2_O, 5.6 mg Na_2_EDTA.2 H_2_O, adjusting to pH 5.8, and 8 g of agar if made for solidification. The hormonal composition that worked for callus development were the variant MS3 for the white line containing 0.2 mg of naphtalene acetic acid and 1 mg kinetin, and MS6 for the red line containing 1mg of naphtalene acetic acid and 0.2 mg kinetin. The obtained calli were kept in their respective solid MS media at room temperature in darkness. Calli were subcultured every 30 days. Cell suspensions were initiated by placing 1.25 g of calli in 100 ml flasks with 25 ml of either liquid MS3 or MS6 media prepared without agar and grown in darkness at 25 °C under orbital shaking at 110 rpm. The volume of the suspension was gradually upscaled to 1L flasks containing 400mL of suspension and then maintained by subculturing every 14 days, by doubling the volume of the suspension adding fresh liquid medium and dividing it between two new sterile flasks.

Elicitation treatments with MBCD and MeJA were carried out in quadruplicate as previously described for grapevine cells (Lijavetzky et al., 2008), with slight modifications. Fresh cells were prepared in an aseptic environment by filtration of a 14-day old *M. alba* cell suspension through a sterile glass filter (90-150 μm pore size) followed by washing with sterile water under gentle vacuum. A weighted amount of cells was transferred into shaking flasks and suspended in fresh growth medium (4 g/g of cell FW) supplemented with elicitors (50 mM MBCD, 100 μM MeJA or 50 mM MBCD + 100 μM MeJA) and was maintained at 25°C in a continuous rotary shaker (110 rpm) in the dark for up to 120 hours. MBCD was added to the medium before autoclave. MeJA prepared as a stock solution of 47.5mM in ethanol was filter-sterilized through a 0,2 μm PTFE filter before adding to the sterile medium. Each replicate consisted in 8 g of cells and 32 g of elicitor-supplemented MS3 or MS6 liquid medium in 100 ml flasks. Controls and MBCD treatment received the same volume of ethanol than MeJA and MBCD-MeJA treatments (2.1 uL/mL cell suspension). Each flask was stored in darkness. The total number of samples was 24, corresponding to a single time point (5 days after elicitation) and three biological replicates per condition (control, MBCD, MeJA, MBCD-MeJA). After incubation, each sample was individually filtered to separate the cells from the media.

Four 14-day old cell suspension flasks grown in 1 L flasks (having around 500 ml of cell suspension) were mixed and filtered using a sterile glass filter (90-150 μm pore size) and a vacuum pump, to separate the cells from the growing media. Each replicate consisted in eight grams of cells and 32 g of liquid media in 100 ml flasks. Control cells were grown in MS6 (for pigmented-fruited cultivar) or MS3 (white-fruited cultivar) media while elicited cells were grown in MS6/MS3 media prepared with MBCD at a final concentration of 50 mM prior to autoclave. Ethanol-diluted MeJA was added after media sterilization. MeJA was filter-sterilized with a 0,2 μm PTFE filter, and added to the MS3 + MBCD media reaching a final concentration of 100 μM (84 μl of a 47,5 mM MeJA stock). Each control flask received 84 μl of 100% ethanol. Cells were kept in orbital agitation at 110 rpm at 24°C. Each flask was protected from light using aluminum foil.

The total number of samples in the time-series experiment was 48, corresponding to six time points (30 min, 6, 12, 24, 48, and 72h after elicitation) and a total of four biological replicates per condition (control and MBCD-MeJA). After the incubation time, each sample was individually filtered to separate the cells from the medium. Media was collected while cells were rinsed with cold sterile water to remove the excess of medium and filtered again. The weight of both cells and media was recorded and samples were quickly frozen in liquid nitrogen and then kept at -80 °C. Cell samples were lyophilized for 48 h, then stored at -80 °C until metabolite, protein or RNA extraction. Media was stored at -20 °C for subsequent metabolic analysis.

### Extraction and LC-MS quantification of stilbenoids

For extraction of fresh or frozen material, two grams of cells were mixed with 4.6 ml of HPLC grade methanol, vortexed and shaken at 1800 rpm for 75 minutes. The remaining debris was filtered using a pasteur pipette and cotton wool. For lyophilized material 10 mg of dry cells were extracted using 1 ml of 80% methanol and incubated at 220 rpm and 4 °C overnight. Extracts were centrifuged at 14.000 x g for 10 minutes and the supernatant was collected for LC-MS analysis (Hurtado-Gaitán et al., 2017). For extracellular stilbenoids, an aliquot of 1 ml of filtered cell-free medium was extracted using 250 μl of HPLC grade ethyl acetate, vortexed, agitated at 1800 rpm for 2 minutes and centrifuged at maximum speed for 5 minutes. The organic phase was collected in a separate tube and a second round of ethyl acetate extraction was conducted. The total pooled volume of organic phase was dried using a SpeedVac at room temperature. The dried residue was resuspended in 500 μl of 80% MS quality methanol centrifuged at 14.000 x g for 10 minutes and the supernatant collected for LC-MS analysis. LC-MS analyses were performed in an Agilent 6490 Triple Quadrupole mass spectrometer coupled with an Agilent 1290 Infinity UHPLC as described in Hurtado-Gaitán et al. (2017), in addition to the MRM methods already developed for quantitative analysis of *Vitis vinifera*’s main stilbenoids (pterostilbene, resveratrol, piceatannol, viniferin and piceid). Here we developed methods for two new stilbenes present in *Morus* species, namely oxyresveratrol and mulberroside A using authentic analytical standards. The in-source fragmentation phenomenon has already been described for piceid, the glycosylated form of resveratrol. Here, we also observed it for mulberroside A, the diglycosylated form of oxyresveratrol, which allows to detect peaks of the aglycone compound formed in the ion source at the retention time of the glycosylated counterpart.

### GC-MS analyses of cell suspensions for umbelliferone detection

The GC/MS analyses were carried out in an Agilent 6890N gas chromatograph equipped with a 30 m × 0.25 mm ID, 0.25 µm film thickness Agilent 19091S-433HP-5MS 5% Phenyl Methyl Silox column, coupled to an Agilent 5973N mass spectrometer as reported in Martínez-Márquez et al. (2018). The column was operated with hydrogen as the carrier gas (1 mL/min) at an initial temperature of 40 °C, which was then raised at a rate of 12 °C/min to 290 °C, held for 6 min, and then raised at 20 °C/min to 320 °C and held at this temperature for 10 min. The injector was kept at 250 °C. Mass spectra were recorded from m/z 33 to 250 at 70 eV. Chromatographic peaks were first identified searching against the NIST Standard Reference Database NIST11.L and then by the retention time and mass spectrum of the authentic compounds.

### Microscopy

*Morus* cells were imaged on a confocal microscopy (Leica TCS SP2; Leica Microsystems, Wetzlar, Germany). Cell suspensions were placed onto microscope slides in MS3 and bright field images were taken in a single-plane of 1 μm. Brightness and contrast were adjusted by Adobe Photoshop 7.0.

### RNA extraction and sequencing

The Spectrum Plant Total RNA kit (Sigma-Aldrich, SKU: STRN250-1KT) was used to extract total RNA for Illumina sequencing from 34 lyophilized cell samples collected during the time-course experiment. For PacBio sequencing, a four-sample pool of different elicited samples was generated. DNAse-treated RNA was monitored for its integrity using an Agilent Technologies 2100 Bioanalyzer defining high quality RNA if the RNA Integrity Number (RIN) value was greater than or equal to 7. For PacBio sequencing, cDNA synthesis was performed using Takara SMARTer PCR cDNA Synthesis Kit for ISO-seq. Molecules under 4Kb were captured for library preparation. The quality of the library was assessed with an Agilent Technologies 2100 Bioanalyzer and QUBIT. Long read sequencing was performed with a PacBio SMRT Cell Seq - RSII (P6-C4). For Illumina sequencing, the TruSeq Stranded mRNA Library Construction kit was used to generate libraries. The quality of the libraries was assessed with an Agilent Technologies 2100 Bioanalyzer and QUBIT. Finally, short read sequencing was performed on an Illumina NovaSeq 6000 producing 150bp paired-end reads.

### Genome re-annotation pipeline

A comprehensive method was implemented to i) revise the protein-coding genes annotated by Jiao et al. (2020) in *M. alba*, evaluate their structure, and reannotate if necessary, and ii) generate new gene models. Supplementary Figure 12 contains a comprehensive compilation of datasets employed, along with a thorough workflow of each step in the bioinformatic pipeline (see Supplementary Table 5 for the parameters used).

MAKER v2 (Campbell et al., 2014) was used to create a new structural annotation integrating data from *de novo* transcriptomes (based on Illumina and PacBio reads obtained in this study), *ab initio* gene prediction and database gene models. The *de novo* transcriptome was constructed using Trinity (Grabherr et al., 2011) in strand-specific mode, utilizing both the Illumina and PacBio evidence. Illumina reads were trimmed using fastp (S. Chen et al., 2018). PacBio sequencing errors were corrected using LoRDEC (Salmela & Rivals, 2014) and Illumina reads. Filtering involved using CD-HIT to retrieve the most representative sequences, Kallisto (Bray et al., 2016) to remove misassembled transcripts while keeping only expressed ones (TPM > 0), and TransDecoder (https://github.com/TransDecoder/TransDecoder) to keep only the protein-coding transcripts. For the *ab initio* prediction of genes, GeneMark software (Besemer & Borodovsky, 2005) was trained using introns annotated by aligning cell culture Illumina reads with STAR (Dobin et al., 2013) on the *M. alba* genome assembly. The Augustus software (Stanke et al., 2006) was also employed, trained with 2,117 genes from the previous annotation. Finally, homology evidence was generated using Moraceae proteins downloaded from the NCBI clustered using CD-HIT (Fu et al., 2012) to reduce redundancy. These proteins were then mapped using BLASTx in MAKER after masking the genome. Masked repetitive regions were interpreted by MAKER using RepeatMasker (version 4.1.2) (N. Chen, 2004) and Dfam database (release 3.5) sequences to locate both previously known and novel transposable elements.

In addition, Minimap2 (H. Li, 2018) and StringTie (Pertea et al., 2015) were used to generate a further annotation file based just on PacBio data. The three annotations; v0 official annotation in Jiao et al. 2020, MAKER-derived, and StringTie-derived, were merged to create a file containing all the possible gene structures.

In order to reduce redundancy in the final annotation, a modified version of TransDecoder, customized to output only complete protein ORFs, was used to filter and retrieve the best possible gene structure for each gene of the merged file. A single protein was then chosen for each transcript. The best protein was selected using Diamond (Buchfink et al., 2015) and interproscan (Paysan-Lafosse et al., 2023) based on the hit scores from a series of databases, prioritized in the following order: Reviewed Viridiplantae proteins from Uniprot with 90% identity, Viridiplantae proteins from NCBI, Viridiplantae proteins from Uniprot with 90% identity, and InterPro domains. Reviewed Uniprot scores were considered the most reliable and subsequent databases were only considered in case of hit score draws. Overlapped transcripts with the same BLAST hit, as well as those with more than 80% overlap, were filtered out to reduce redundancy, aiming to have one transcript per gene. The best transcript was chosen according to the same database priority used above. If several transcripts for a gene had the same hit score (based on E-value and Bit-score), the transcript was then chosen by highest FPKM expression or, if this information was unavailable, by longest transcript.

The annotation was evaluated at a protein level against the annotated proteins Jiao et al. (2020) using BUSCO over the eudicots_odb10 dataset (Simão et al., 2015). We introduced a new gene ID that contains information about the species, cultivar, assembly and annotation versions, with the fixed prefix of “Mal_HE_154X_1”.

### Functional annotation

The functional annotation of genes was performed using eggNOG-mapper (Cantalapiedra et al., 2021), KAAS (Moriya et al., 2007), and Mercator4 (Schwacke et al., 2019) web applications for GO, KEGG, MapMan ontologies, respectively. All pipelines were run with default settings, except for KAAS, where some of the best annotated plant species were selected: *Arabidopsis thaliana* (ath), *Zea mays* (zma), *Vitis vinifera* (vvi), *Glycine max* (gmx), and *Solanum lycopersicum* (sly).

### DEA and WGCNA

Illumina reads were trimmed using Fastp software (S. Chen et al., 2018), followed by mapping using STAR (Dobin et al., 2013). Raw counts were computed using FeatureCounts (He et al., 2013), where the new annotated gene models were used. Normalized counts were used for a principal component analysis (PCA) (Supplementary Figure 13A). Differential expression analysis was performed using the LIMMA R package (Ritchie et al., 2015), while FPKM normalization was conducted using the DESeq2 package (Love et al., 2014). Genes were considered differentially expressed when the adjusted p-value was lower than 0.05. For co-expression analysis, we used the weighted gene correlation network analysis (WGCNA) R package (Langfelder & Horvath, 2008), with FPKM values for all samples filtered by expression using the filterByExpr function of edgeR (Robinson et al., 2010). We imported the data for resveratrol and oxyresveratrol synthesis to perform module-trait correlations. Additionally, enrichment analysis for Mapman, KEGG, and GO for different modules was performed using the gprofiler2 R package (Kolberg et al., 2020).

### Phylogenetic analysis

Sequences were aligned using MAFFT (Katoh, 2002). Obtained alignments were used to build maximum-likelihood trees with IQ-TREE software (Nguyen et al., 2015) and visualized with FigTree. The bootstrap consensus tree was inferred from 1000 replicates.

### Protein extraction and identification

Lyophilized cells were first washed according to (Lücker et al., 2009) with modifications. All subsequent steps were carried out on ice and centrifugations were performed at 4°C. Throughout the procedure, each wash was followed by centrifugation for 15 min at 15300 x g. Briefly, 80-130 mg lyophilized cells were washed in ethyl-acetate:ethanol 1:2 (v/v) several times, until the supernatant was colorless. The pellet obtained was washed twice with chilled acetone and twice with 10% trichloroacetic acid (TCA) (w/v) in acetone - 20°C. This was followed by three times washing with 10% aqueous TCA (v/v). Finally, pellets were washed twice with 80% chilled acetone (v/v) and left to dry at 4°C.

Total proteins were extracted from the above washed tissue according to Hurkman & Tanaka (1986) with modifications. Briefly, all washed tissue was homogenized in 1ml extraction buffer pH 7.5 (0.7 M sucrose, 0.1 M KCl, 0.5 M Tris, 50 mM EDTA, 1% PVPP, 1% DTT, 1% deoxycholate and a cocktail of protease inhibitors containing 4-(2-aminoethyl) benzenesulfonyl fluoride (AEBSF), E-64, bestatin, leupeptin, aprotinin, and sodium EDTA (Sigma-Aldrich)) and incubated during 30 min with frequent vortexing. An equal volume of Tris-saturated Phenol pH 7.5 was added and the mixture was incubated during 30 min with vortex every 5 min. Separation of the phases was achieved by centrifugation at 15000 x *g* for 40 min. The upper phenol phase was recovered and the aqueous phase was re-extracted as mentioned above with 1ml Tris-saturated Phenol pH 7.5. Both phenol phases were pooled and washed twice with an equal volume of washing buffer at pH 7.0 containing 0.7 M sucrose, 0.1 M KCl, 0.5 M Tris, 50 mM EDTA, 1% DTT and protease inhibitors as above. The recovered upper phenol phase was precipitated for 24h with 5 volumes of 0.1 M ammonium acetate in methanol. The precipitate obtained was washed three times in cold 0.1 M ammonium acetate in methanol and twice in chilled 80 % acetone (v/v) allowed to dry, and solubilized in fresh 6M urea. Protein was quantified by RC DC protein assay (BIO-RAD) based on the modified Lowry protein assay method (Raghupathi & Diwan, 1994).

### Label-free proteomic analysis

A time series proteomic experiment was carried out using quadruplicates of the whole cell extracts from the mulberry cell suspensions control and treated with 50 mM MBCD+0.1 mM MeJA elicitors for 24, 48 and 72 hours. Trypsin protein digestion and peptide clean-up was performed as described in Esteve-Sánchez et al. (2020). Thirty micrograms of the desalted peptide digests were injected directly onto a reverse phase Agilent AdvanceBio Peptide mapping column (2.1 mm × 250 mm, 2.7 μm particle size) mounted in an Agilent 1290 Infinity UHPLC coupled through an Agilent Jet Stream® interface to an Agilent 6550 iFunnel Q-TOF mass spectrometer (Agilent Technologies) system. Peptides were separated at 50 °C using a 140 min linear gradient of 3-40 % ACN in 0.1 % formic acid at 0.400 mL/min flow rate. Source parameters employed gas temp (250 °C), drying gas (14 L/min), nebulizer (35 psi), sheath gas temp (250°C), sheath gas flow (11 L/min), capillary voltage (3,500 V), fragmentor (360 V). The data were acquired in positive-ion mode with Agilent MassHunter Workstation Software, LC/MS Data Acquisition B.08.00 (Build 8.00.8058.0). The mass spectrometer was operated in high sensitivity mode and MS and MS/MS data were acquired in Auto MS/MS mode whereby the 20 most intense parent ions (charge states from 2 to 5) within 300 to 1,700 m/z mass range above a threshold of 1,000 counts were selected for MS/MS analysis. MS/MS spectra (50 - 1,700 m/z) were collected with the quadrupole set to “narrow” resolution and were acquired until 25,000 total counts were collected or for a maximum accumulation time of 333 ms.

Each MS/MS spectra was preprocessed with the extraction tool of Spectrum Mill Proteomics Workbench (Agilent) to obtain a peak list and to improve the spectral quality by merging MS/MS spectra with the same precursor (Δm/z<1.4 Da and chromatographic Δt < 15s). The reduced dataset was searched against the *M. alba* reannotated protein database (Mal_HE_154X_1) and contaminant proteins in the identity mode with the MS/MS search tool of Spectrum Mill Proteomics Workbench and with the following settings: trypsin, up to 2 missed cleavages, carbamidomethylation of Cys as fixed modifications, oxidation of Met as variable modification and mass tolerance of 20ppm for precursor and 50ppm for product ions. Peptide hits were filtered for score ≥ 6 and percent scored peak intensity (%SPI) ≥ 60.

The LC-MS raw files were imported into Progenesis QI for Proteomics (Nonlinear Dynamics) v4.0 label-free analysis software. Quantification was done on the basis of MS1 intensity. The data file that yielded most features (peaks) was used as reference to align the retention time of all other chromatographic runs and to normalize MS feature signal intensity (peak area). Correction for experimental variations was done by calculating the robust distribution of all ratios (log(ratio)). The MS features were filtered to include only features with charge state from two to five. After defining the experimental design as “between subjects” mode samples were clustered according to the experimental groups (C24, C48, C72, MBCD-MeJA24, MBCD-MeJA48 and MBCD-MeJA72) and the average intensity ratios of the matched features across the experimental groups as well as the p-value of one-way ANOVA were automatically calculated. To identify the proteins from which detected features come from, the filtered SpectrumMill peptide hit files were imported into Progenesis QIp. Peptide assignment conflicts were resolved in favor of the highest scoring ones (or left unresolved in cases of equal scores and sequence) and the inferred protein list was filtered by score ≥ 15. Protein abundance was automatically calculated by the Hi-3 method as described by Silva et al. (2006) implemented in the Progenesis QI for proteomics. Differential protein abundance across experimental groups was assessed by using the advanced statistical tools implemented in Progenesis QIp including ANOVA, hierarchical clustering and power analysis. Detected proteins with ANOVA significance p>=0.05 (410 proteins) were used for a principal component analysis (PCA) (Supplementary Figure 13B).

### Protein-RNA correlation analysis

The Spearman’s rank correlation coefficient (ρ) of each mRNA-protein pair was used for correlating transcriptome and proteome levels. Transcriptomics and proteomics data were evaluated by means of Weighted Gene Co-expression Network Analysis (WGCNA) (Langfelder & Horvath, 2008). FPKMs and normalized protein abundances were separately transformed into z-scores, and WGCNA was performed with a soft-power of 6 signed network for DEG-DAP mRNA-protein pairs. Modules were defined by dynamic tree cut with a minimum size of 10 and deep split of 4.

### Cloning of C2’H cDNA and construction of the binary vector

First-strand cDNA was synthesized from total RNA using a cDNA synthesis kit (NZY First-Strand cDNA Synthesis Kit, (NZYTech) according to the manufacturer’s instructions. *C2’H1* and *C2’H4* coding regions (Supplementary File 1) were PCR amplified from a pool of elicited cell RNAs from different time-points (primers used are shown in Supplementary Table 6). The amplification reactions consisted of a polymerase activation cycle of 2 minutes at 95 °C, 40 cycles of denaturation (20 seconds at 95 °C), annealing (10 seconds at a selected temperature) and extension (20 seconds at 70 °C), a final extension cycle of 10 minutes at 70 °C and a final cycle at 4 °C. Annealing temperatures were set by selecting the lowest melting temperature of the primers used. Amplified DNA fragments were cloned into pENTR/D-TOPO plasmids (Invitrogen) as recommended by the manufacturer, and the inserts sequenced by LightRun Tube service (Eurofins, Germany), using Sanger Sequencing. Correct inserts were transferred into Gateway-compatible vector pB2GW7 under the CaMV35S promoter control using an LR clonase reaction (Invitrogen, Thermo Fisher) carried out according to the manufacturer’s instructions. This construct contained Spectinomycin selectable marker reporter gene. The *C2’H5* gene was obtained from commercial synthesis (Gene-Script; Piscataway NJ, USA) and also transferred into the Gateway compatible vectors pB2GW7 and pKAN-ALLIGATOR2. The resulting binary vectors (Supplementary Figure 14), plus pJCV52-VviSTS42 and pBINY53-VviSTS48 were transferred into chemically competent *Rhizobium radiobacter* (*Agrobacterium tumefaciens*) strain C58 by by freeze and thaw method and selected in LB plates supplemented with spectinomycin and rifampicin. Colonies where cheeked by PCR.

### Transient expression in *N. benthamiana*

pB2GW7-C2’Hs and pBINY53-VviSTS48 were infiltrated in *N. benthamiana* leaves. Agrobacterium C58C1 harboring the vector pCH32-35S:p19 which expresses the silencing suppressor p19 of tomato bushy stunt virus was also used. All different agrobacteriums where cultured in liquid media to late exponential phase and cells were harvested by centrifugation at 3000g for 15 minutes at room temperature (24 °C). Cell pellets where resuspended in agroifiltration buffer (10 mM morpholinoethanesulphonic “MES” acid-KOH, 10 mM MgCl2 and 150 mM acetosyringone; pH 5.6) and incubated for 3 h at room temperature (24 °C). These cells were mixed in different combinations, with a 1:1 ratio, to a final total agrobacterium OD600 of 1, and then injected into young fully expanded leaves of 4-week-old *Nicotiana benthamiana* plants (Schöb et al., 1997). Additionally, control samples where obtained injecting resveratrol (70 μM in 10 mM MES) 4 days after agroinfiltration and oxyresveratrol (70 μM in 10 mM MES) in no agroinfiltrated leaves. Leaves samples were collected 3 and 6 days after agro-infiltration and stored at -80°C.

### Transient expression of grapevine cells

*A. tumefaciens* harboring constructs were used to transiently transform *Vitis* cell suspensions. Transient transformation experiments were performed as described in Martínez-Márquez et al. (2023). The strains harboring the binary plant vectors pKAN-ALLIGATOR-C2’H1, used alone or mixed with a strain containing pJCV52-VviSTS42 (Hidalgo et al., 2017), were co-cultured in a 1:1 ratio in *Vitis* cell suspensions. At 6 days after Agrobacterium-infection, stilbene content was analyzed.

### Web apps

The genomic database MorusalbaDB was created based on EasyGDB (Fernandez-Pozo & Bombarely, 2022), including the download of the annotation and ontology files, BLAST, genome browser, sequence extraction, and gene search. We also deployed a Web Apollo platform (Lee et al., 2013) to manually curate the gene models identified from the annotation pipeline. The MulViz data visualization platform is hosted under a shiny server to deploy each R based web application.

