## Supplemental Figures for "Biosynthesis of oxyresveratrol in mulberry (*Morus alba* L.) is mediated by a group of p-coumaroyl-CoA 2’-hydroxylases acting upstream of stilbene synthases"

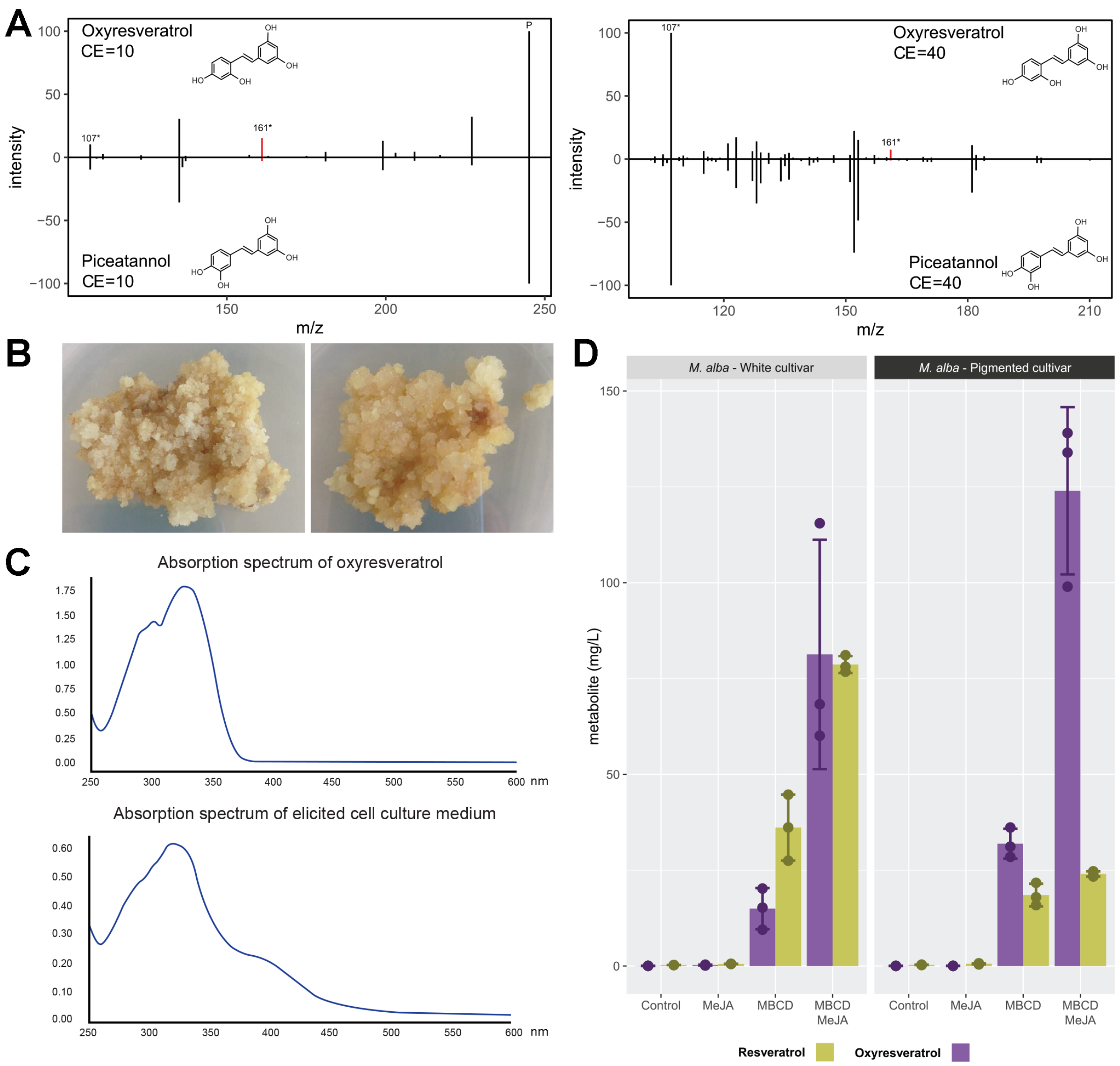


**Supplementary Figure 1** . A) Mirror plot showing mass spectra of oxyresveratrol (top) and piceatannol (bottom) for collision energies (CE) of 10 eV (left) and 40 eV (right). Discriminatory fragment ion (m/z: 161) is highlighted. Precursor ion (P) is not observed at the maximum CE used in this study. B) Calli from white (left) and pigmented (right) cultivars of *Morus alba*. C) Absorption spectrum of oxyresveratrol (top) and elicited cell culture medium at 5 days post treatment (bottom). D) Accumulation of resveratrol and oxyresveratrol in response to single or combined treatment of methyl-β-cyclodextrins and methyl jasmonate (5 days post elicitation).

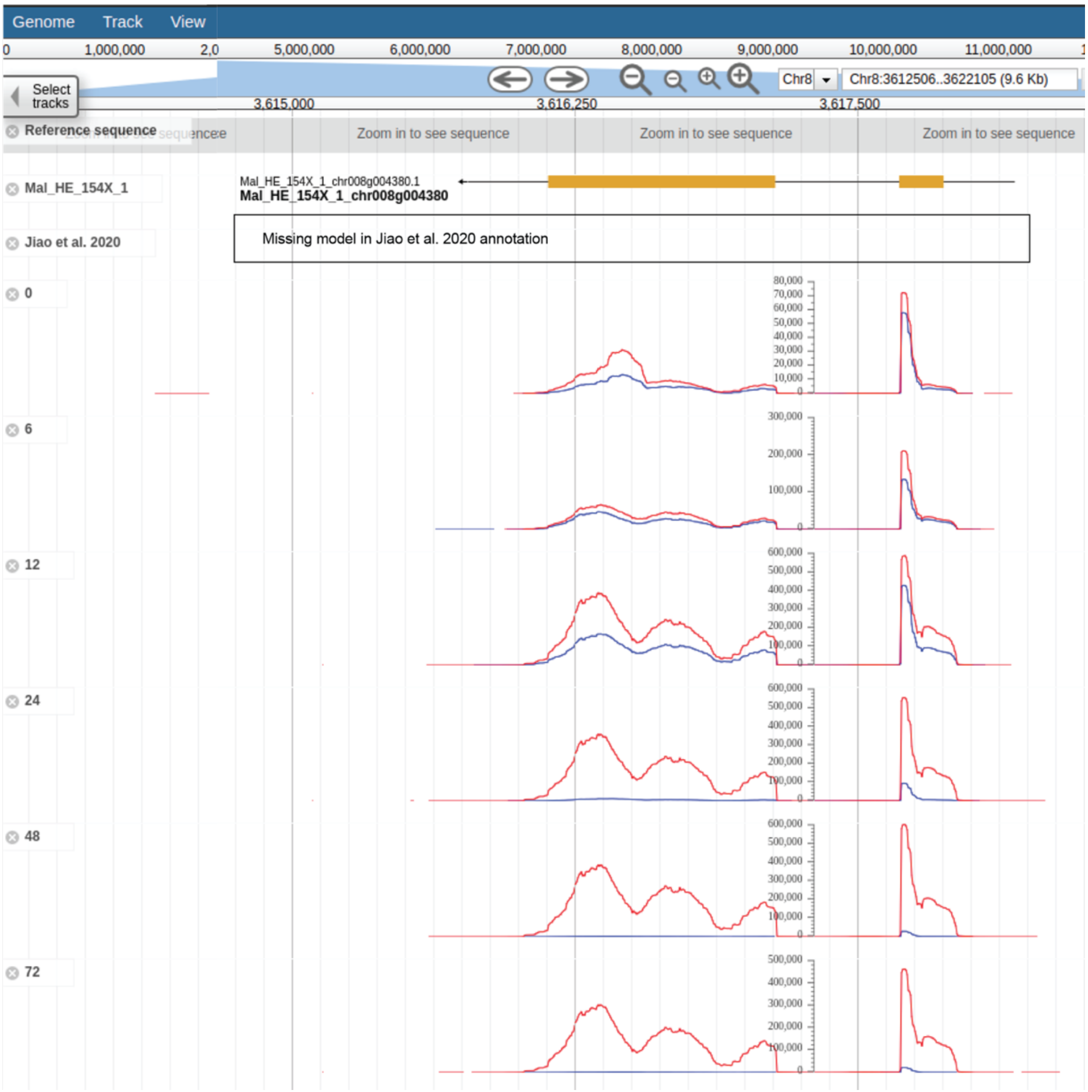


**Supplementary Figure 2. Gene structural annotation of an *STS* gene in the mulberry (Morus alba) genome.** Screenshot of a JBrowser showing the gene model of *STS2,* identified for this first time in this study (the model was not present in the official annotation of (Jiao et al., 2020). Histograms of total aligned counts for *MalSTS2* at different time-points of control (blue) and elicitation (red): 0.5h (30min), 6h, 12h, 24h, 48h and 72h post elicitation.

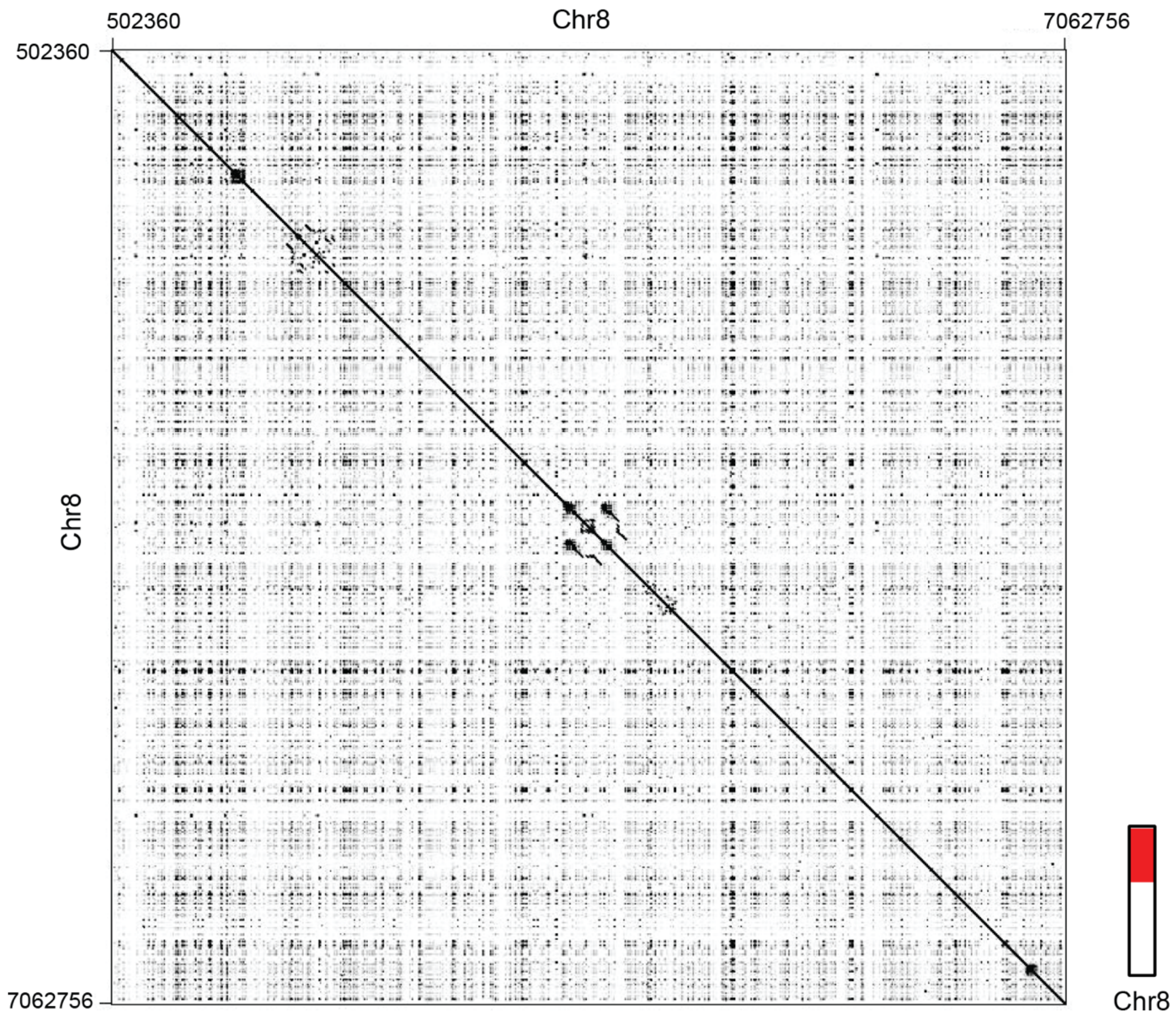


**Supplementary Figure 3. Sequence comparison (dotplot) of a 6.5 Mb genomic region in chromosome 8 of *Morus alba*.** The region is depicted in red at the chromosome level. The center of the plot shows the two *STS* gene clusters shown with greater detail in Figure 2.

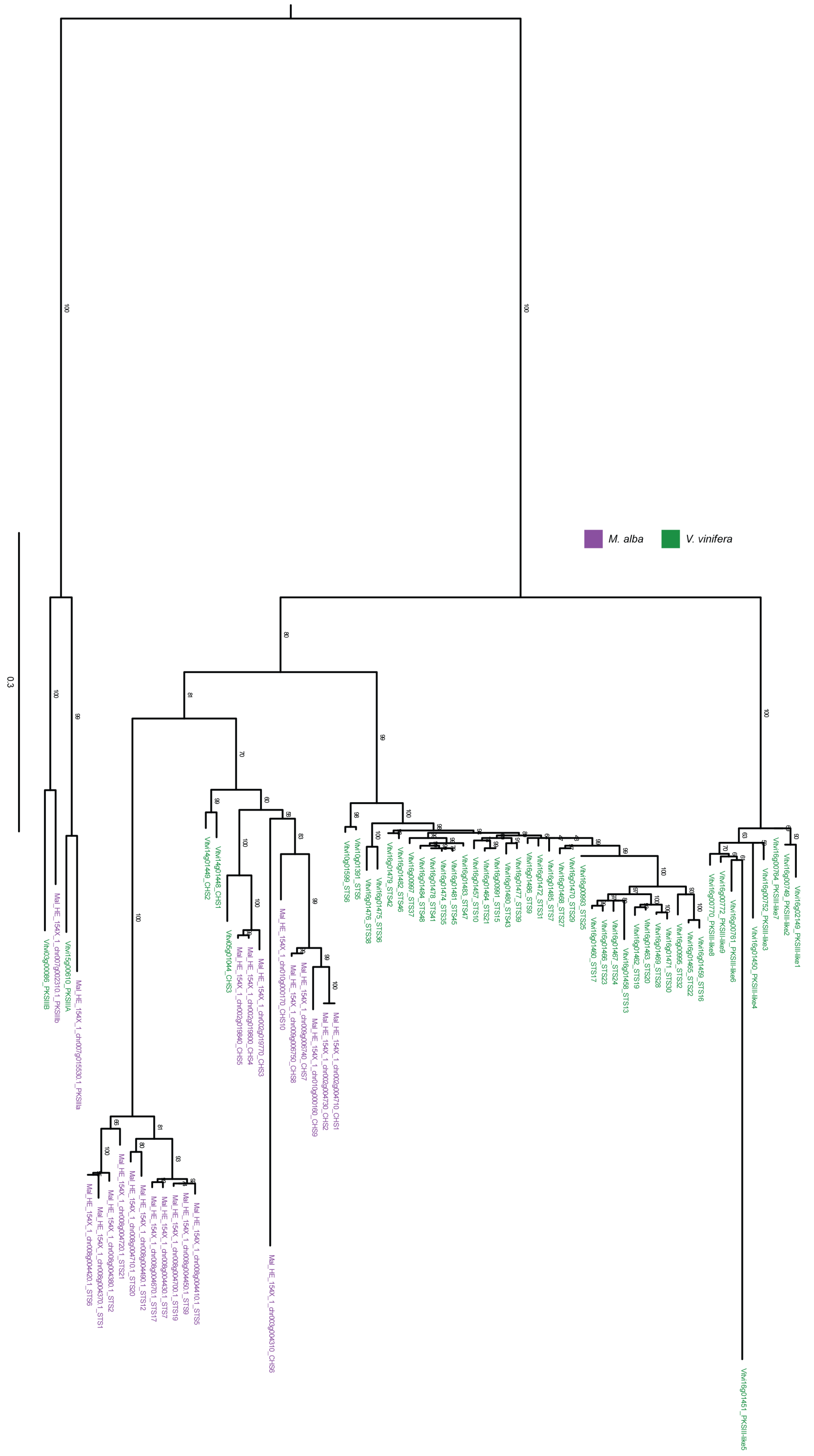


**Supplementary Figure 4. Maximum likelihood phylogeny of the STS, CHS and PKSIII-like protein families.** PKSIIIA and PKSIIIB enzymes from *V. vinifera* were included. The number on top of the branches correspond to the bootstrap values associated with each branch. Scale bar represents substitutions per nucleotide site.

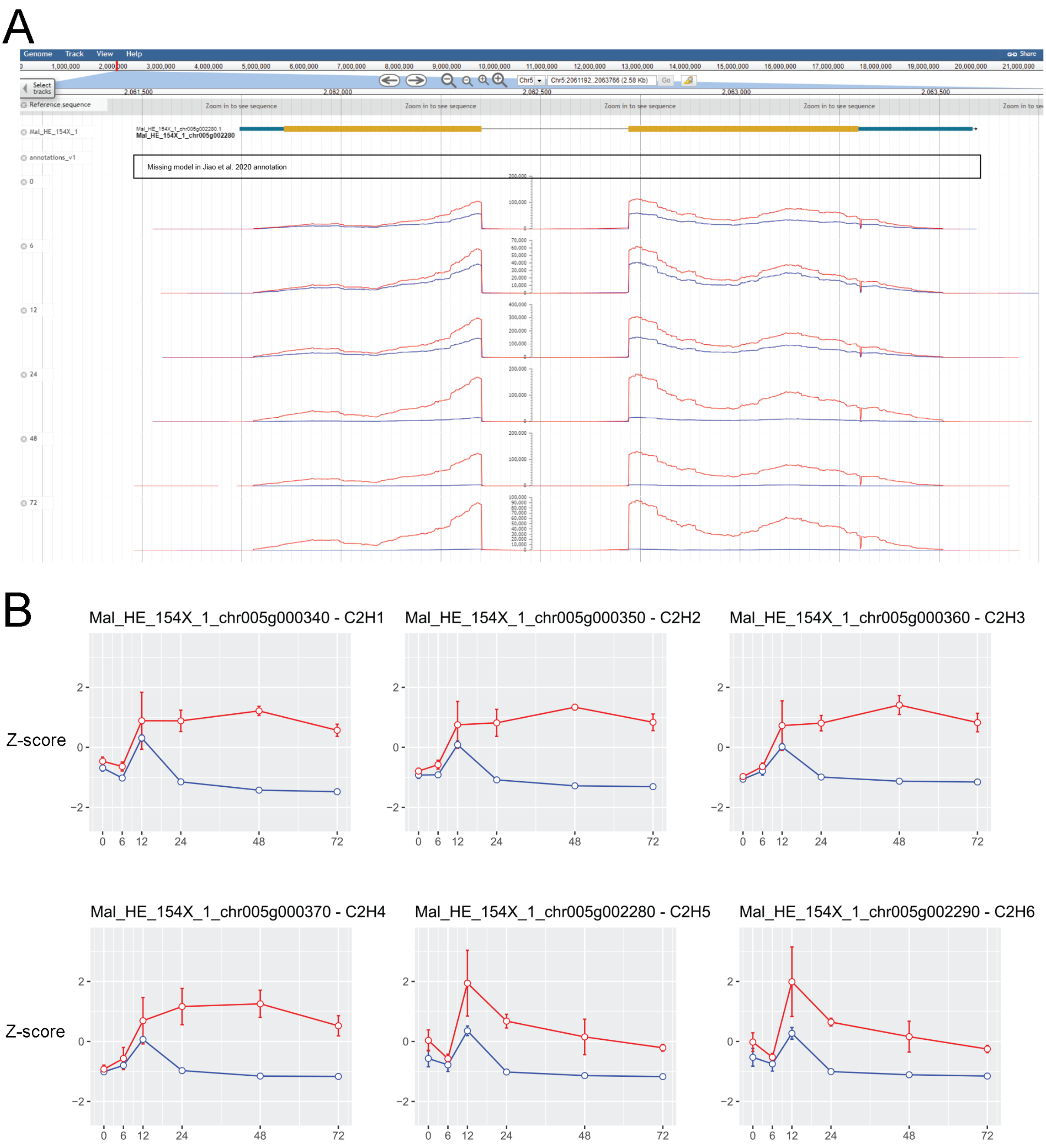


**Supplementary Figure 5. The putative *F6H*/*C2'H* gene family in mulberry**. A) Screenshot of a JBrowser showing the gene model of *C2'H5,* identified for this first time in this study (the model was not present in the official annotation of (Jiao et al., 2020). Histograms of total aligned counts for *C2'H5* at different time-points of control (blue) and elicitation (red): 0.5h (30min), 6h, 12h, 24h, 48h and 72h post elicitation. B) Gene expression of the putative *C2'H* gene family. Expression values were normalized by FPKM and Z-score scaling, independently applied for each gene across the different timepoints of control/elicitation.

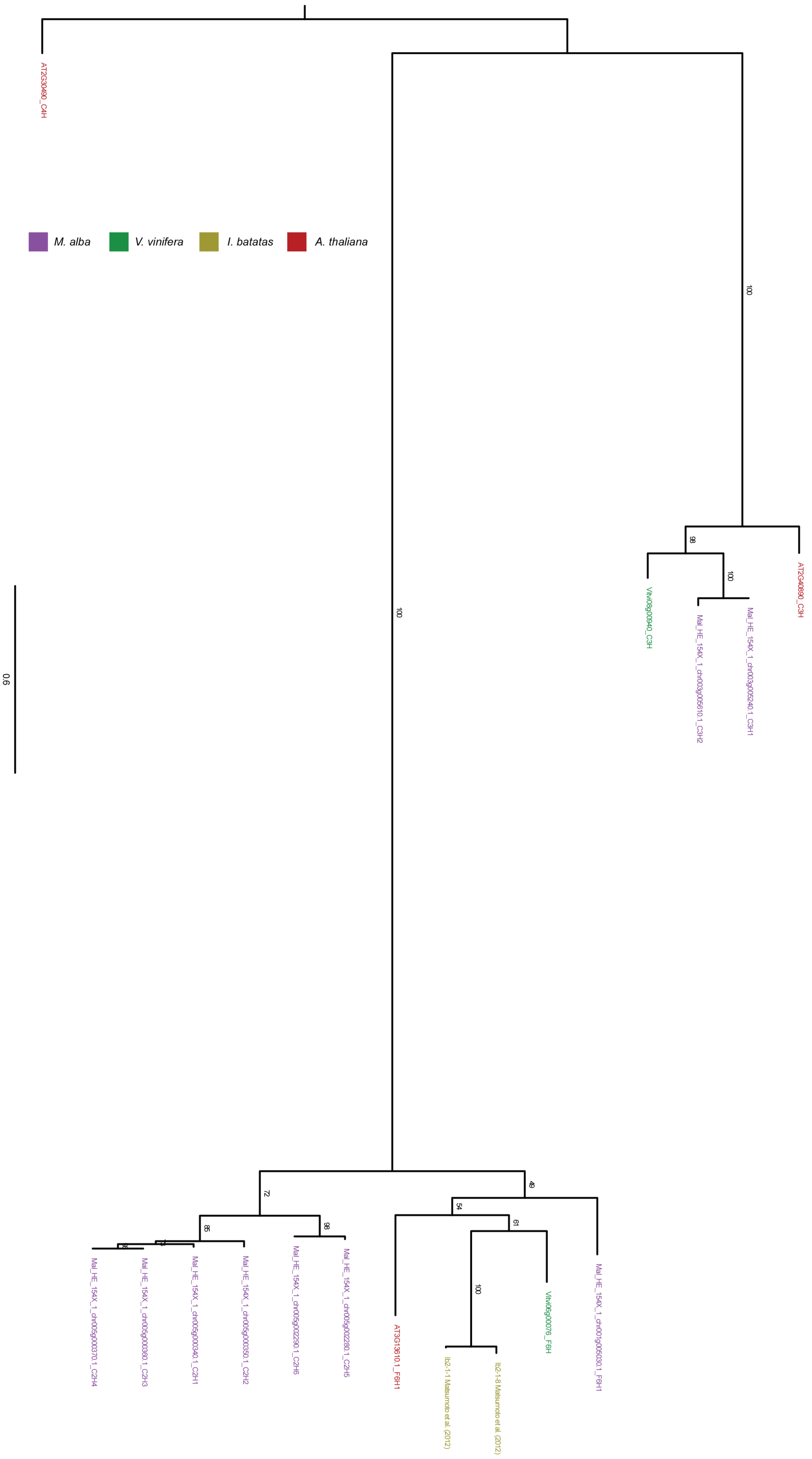


**Supplementary Figure 6.** Maximum likelihood phylogeny of C2’H and C3’H proteins from *Morus alba*, *Vitis vinifera* and *A. thaliana*. C4’Hs proteins were used as an outgroup. The number on top of the branches correspond to the bootstrap values associated with the branch. Scale bar represents substitutions per nucleotide site.

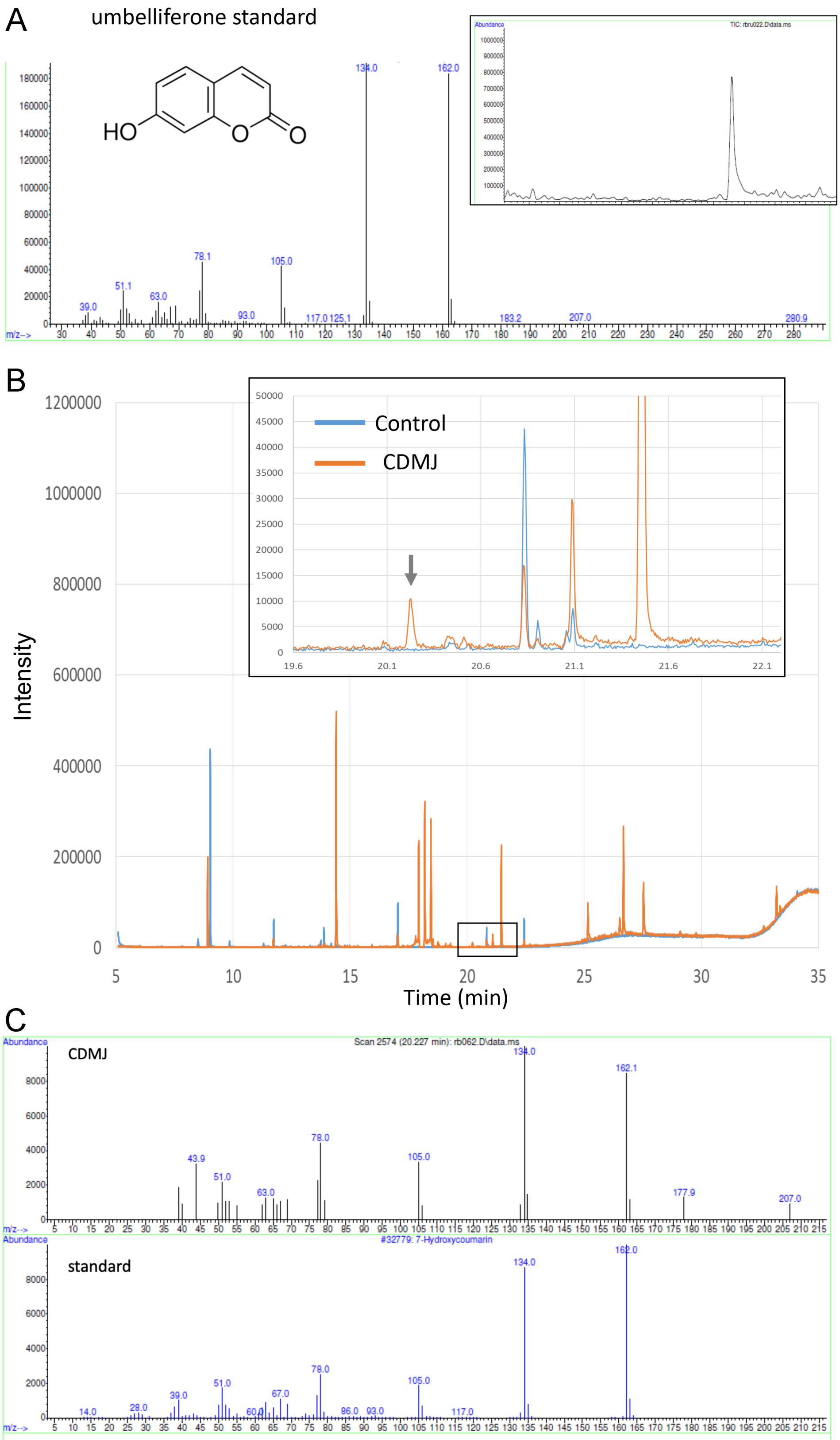


**Supplementary Figure 7. Detection of umbelliferone by GC-MS in elicited mulberry cells.** A) fragmentation spectrum of the analytical standard at the GC base peak. B) Comparison of TIC chromatographic traces in control and elicited conditions. Zoom-in square: arrow points to the differential umbelliferone peak, only present under elicitation. C) Comparison of fragmentation spectra between umbelliferone standard and the elicited differential peak.

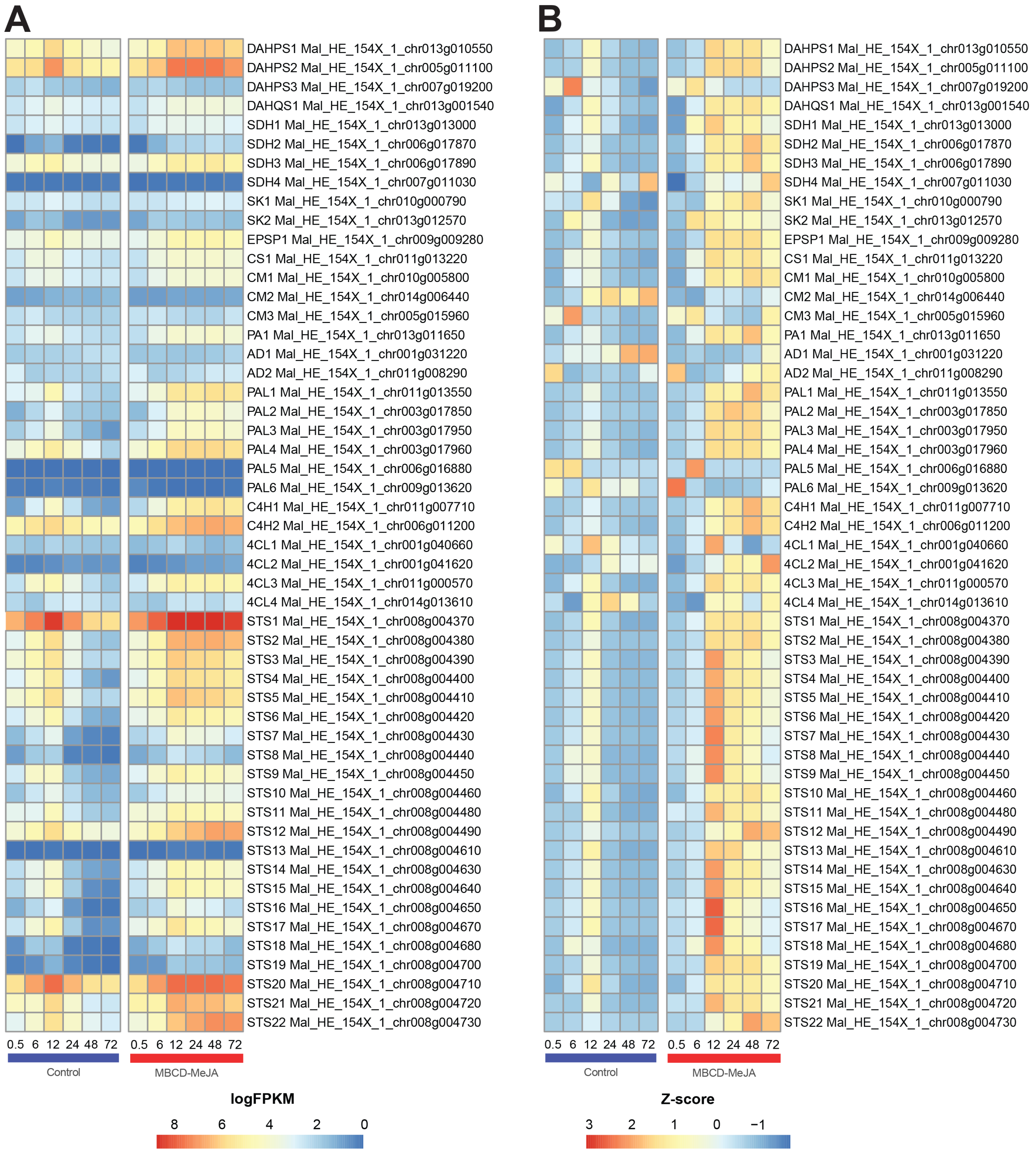


**Supplementary Figure 8. Gene expression changes of the Shikimate and Early phenylpropanoid pathways in response to elicitation in Morus cell suspensions.** A) log-scaled FPKM values. B) Z-score scaling of FPKM normalized values, calculated independently for each gene.

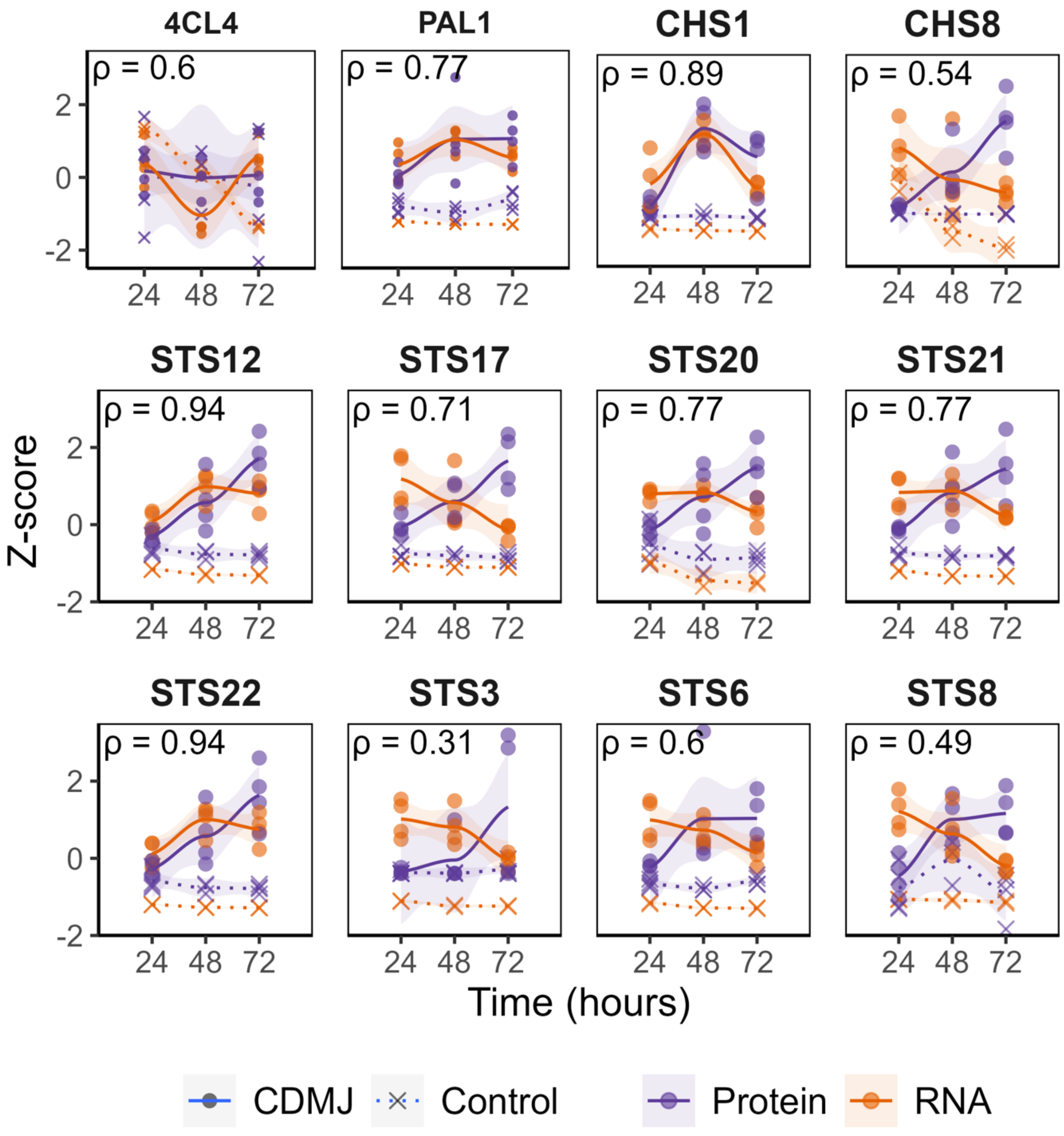


**Supplementary Figure 9. Trajectories of DEG-DAP mRNA-protein pairs at 24, 48 and 72h post elicitation.** Z-scored abundances of RNA and protein levels (Z-scored separately) of selected phenylpropanoid-related enzymes.

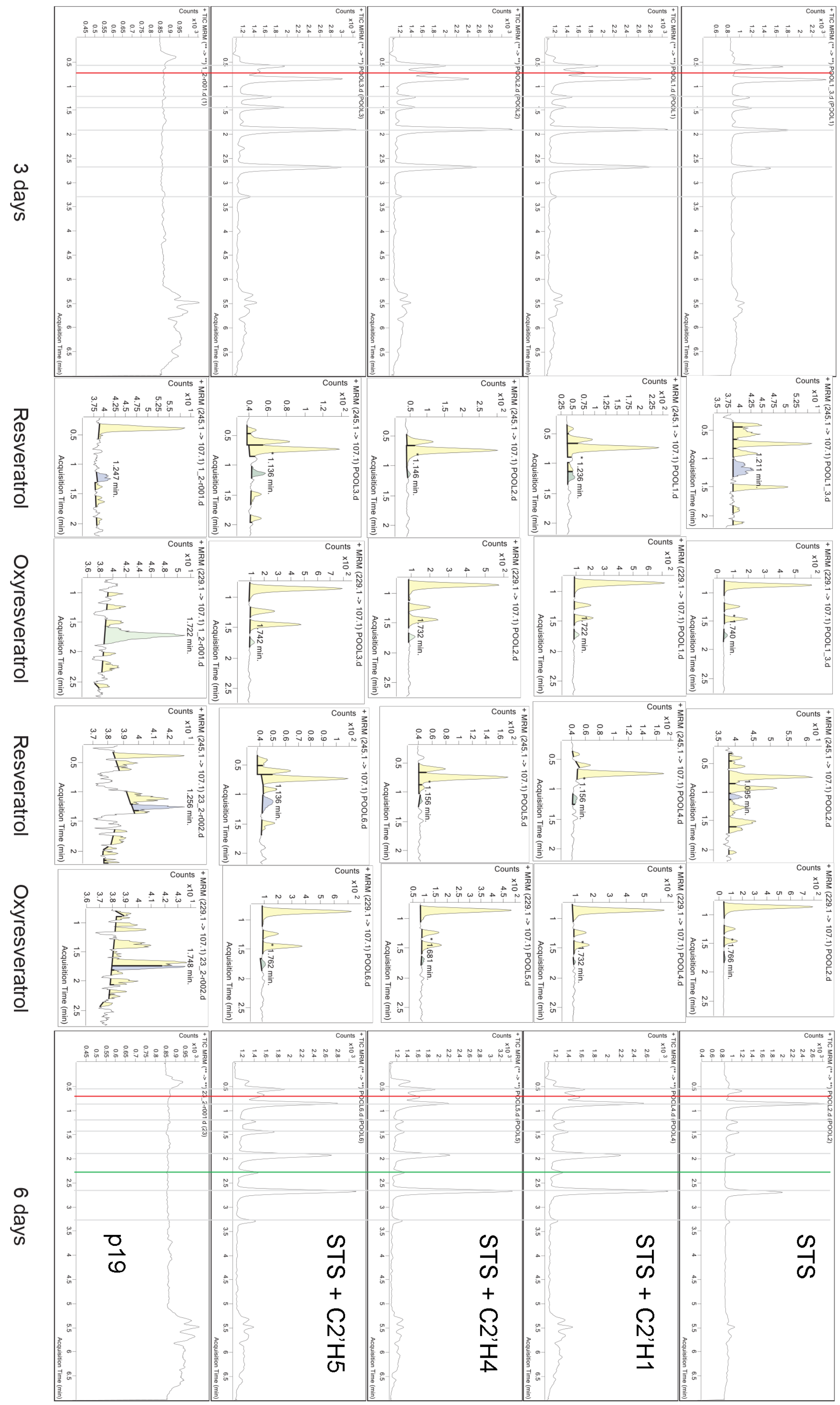


**Supplementary Figure 10.** Representative LC-Mass Spectrometry chromatograms obtained from methanol extracts of agroinfiltrated *Nicotiana benthamiana* leaves. Leaves were co-infiltrated with *Agrobacterium tumefaciens* harboring the P19 helper, and *STS* and/or *C2'H* genes, and left under normal plant growth conditions for 3 and 6 days.

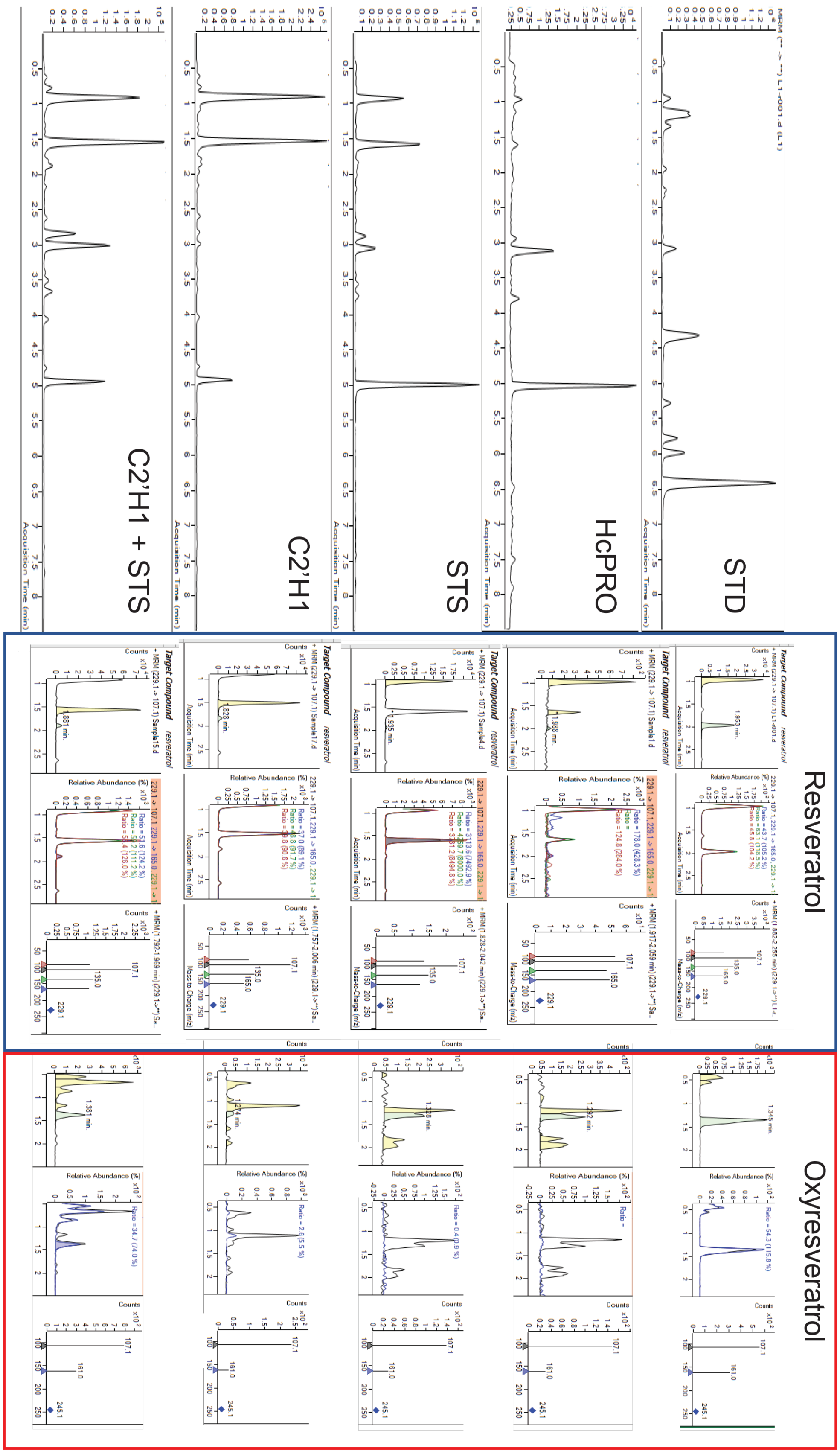


**Supplementary Figure 11.** LC-Mass Spectrometry chromatograms obtained from transiently transformed *Vitis vinifera* cells with *STS* or *C2'H* genes. Samples were taken at 6 days post Agrobacterium co-cultivation.

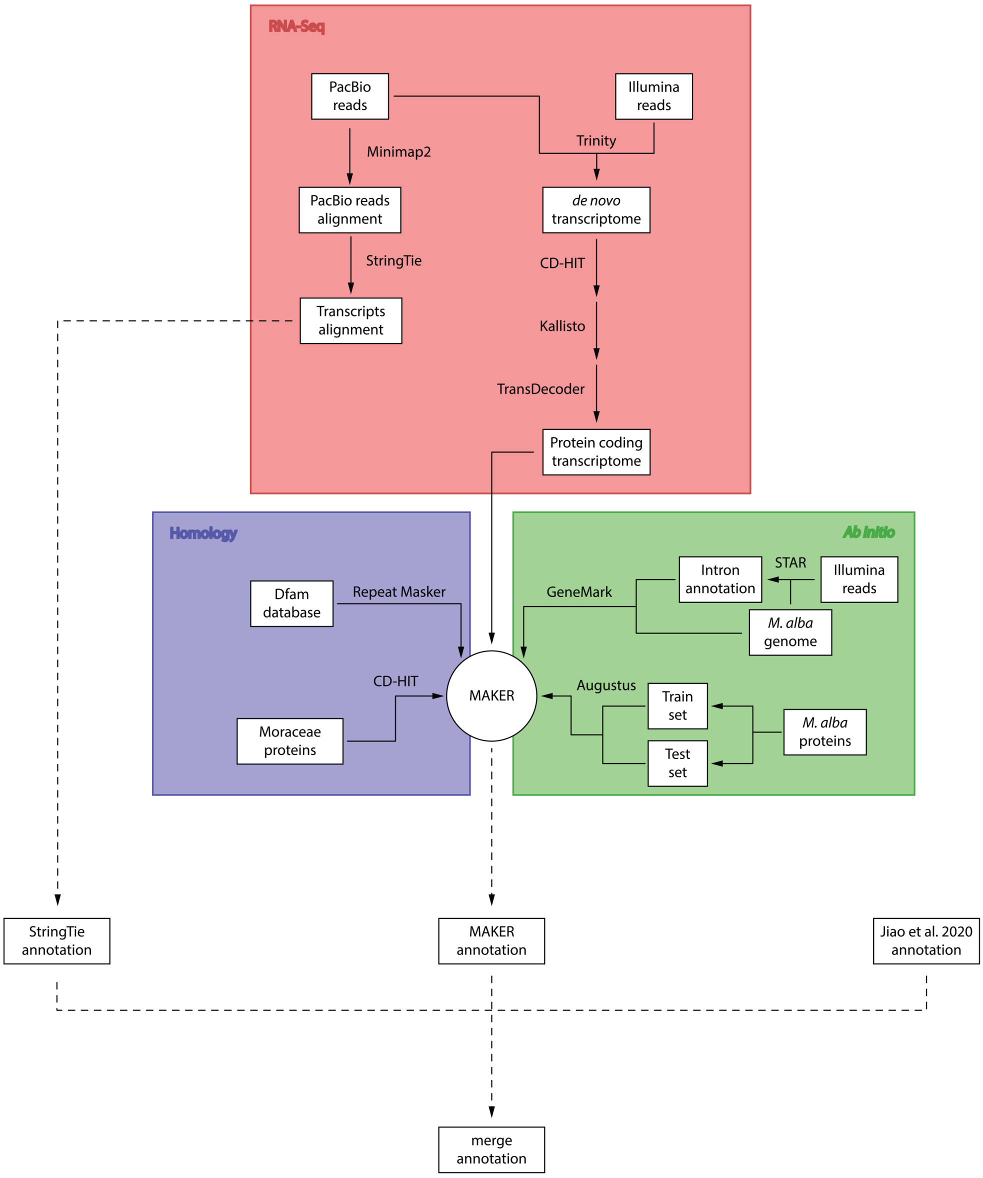


**Supplementary Figure 12. Comprehensive flowchart of the bioinformatics pipeline utilized in the structural reannotation of the *M. alba* genome.** From the RNA-Seq branch (in pink), long-reads from PacBio and short-reads from Illumina were processed to align transcripts and identify protein-coding regions. CD-HIT and Kallisto were applied to refine the *de novo* transcriptome, ending with a TransDecoder step to identify the protein-coding transcriptome. The homology branch (in purple) leverages the Dfam database and RepeatMasker for identifying repetitive elements and employs Moraceae proteins for homology-based evidence generation, after redundancy reduction using CD-HIT. The *ab initio* predictions (in green) use GeneMark and Augustus, trained with a subset of annotated introns and proteins, to predict gene structures directly from the genome sequence. The Illumina reads provide additional support for intron annotation through the STAR aligner. These independent analyses converge into the MAKER annotation engine, which synthesizes evidence from each method to produce a unified and comprehensive annotation of the *M. alba* genome. Dotted lines highlight the flow of data between methods, ultimately leading to a merged final annotation that combines the StringTie annotations with those from MAKER, incorporating both the latest and previously established annotations (referenced by Jiao et al. 2020) for an updated genomic insight.

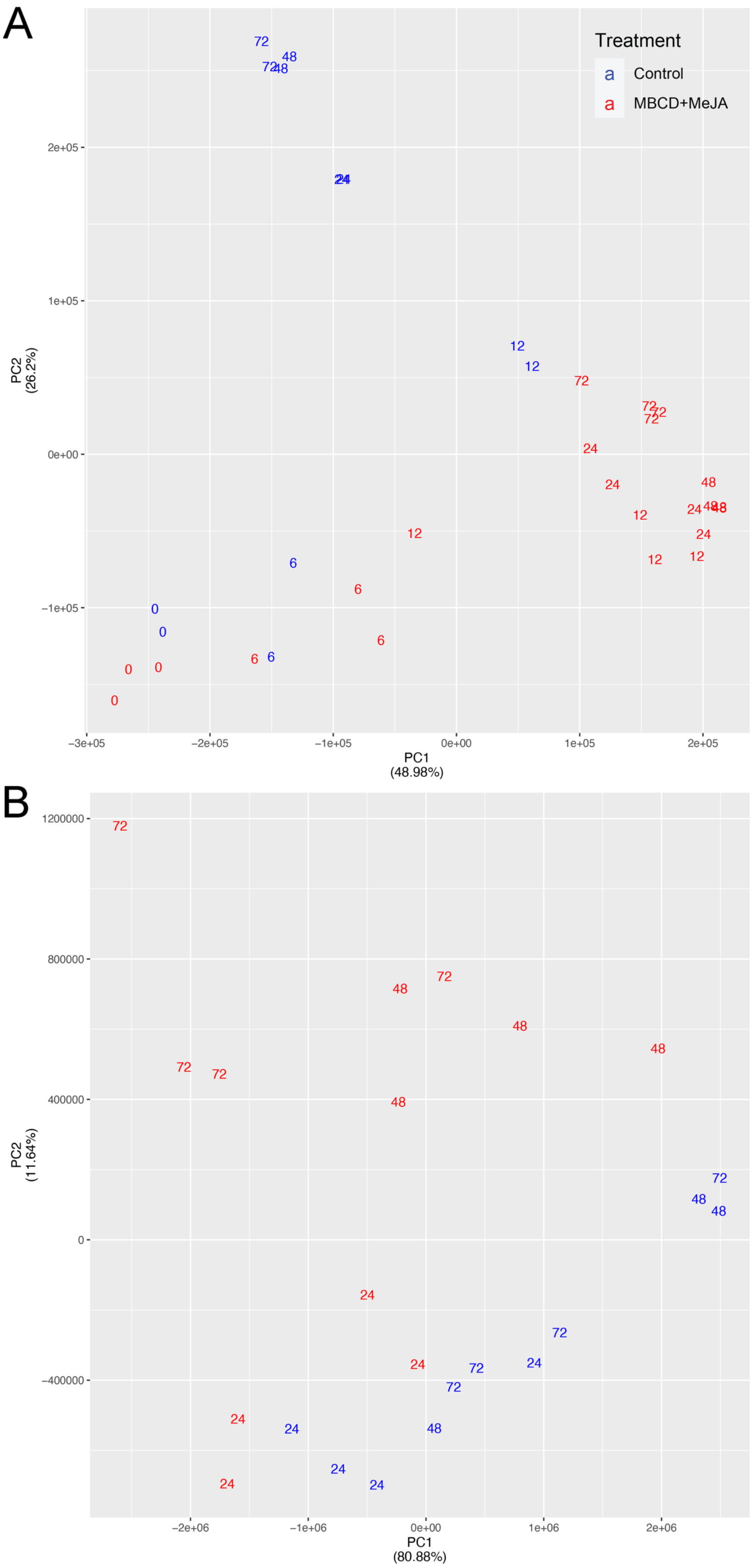


**Supplementary Figure 13.** Transcriptome (A) and proteome (B) two-dimensional score plots (principal component analysis; PCA) of control and MBCD-MeJA elicited samples. The numbers represent the different timepoints (h), with each sample being a biological replicate.

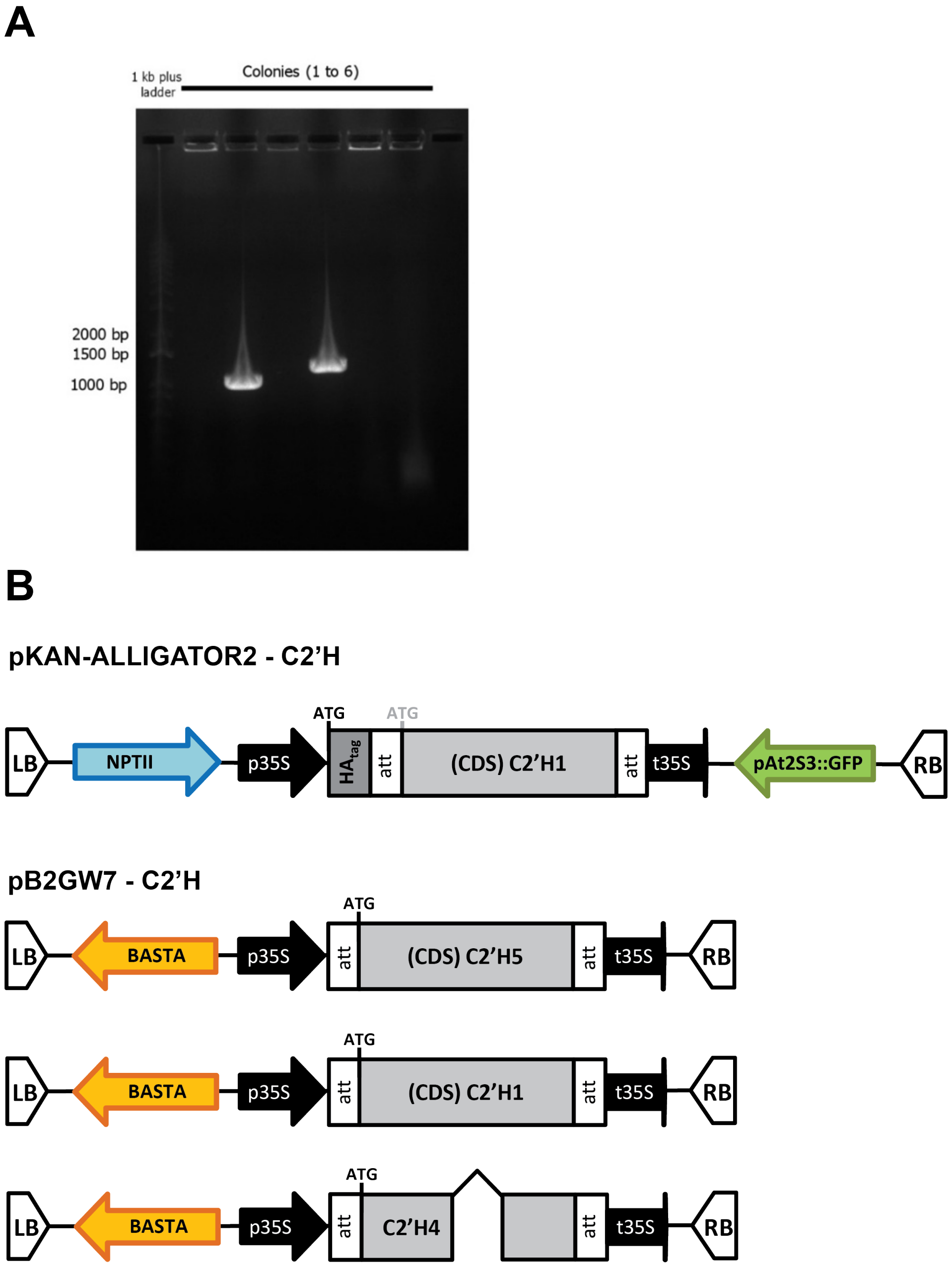


**Supplementary Figure 14.** A) Colony-PCR and 1% agarose gel electrophoresis analysis of pENTR-D/TOPO cloned *C2'H* gene sequences. 1 kb plus ladder (Invitrogen), PCR product of six E. coli colonies: pENTR-D/TOPO-MalC2'H1 (third well) and pENTR-D/TOPO-MalC2'H4 (fifth well). B) Schematic representation of the vectors used in the agrobacteria transient expressions of *Nicotiana benthamiana* (Fig 4F) and Vitis cells (Fig 4G). *NPTII*: neomycin phosphotransferase II; p35S: Cauliflower mosaic virus 35S promoter; t35S: Cauliflower mosaic virus 35 S terminator. pJCV52-VviSTS42 (Hidalgo et al. 2017) and pBINY53-VviSTS48 (Santos-Rosa et al. 2008) were also used in some combinations (constructs not shown).
