## Supplemental File 1 for "Biosynthesis of oxyresveratrol in mulberry (*Morus alba* L.) is mediated by a group of p-coumaroyl-CoA 2’-hydroxylases acting upstream of stilbene synthases"

>FWZ793_39777936_39777936_C2’H1_Mal_HE_154X_1_chr005g000340

Atggcgcctaccaaaaccagtgccgcacaatgggataacgattctgtaactgagtgggtgataaacaaaggcaatggagtgaagggtctctcagaaactggcatcacaaccctcccccatcagtatatccagccagtggaagagaggacgatgagtaaggtcttggtcggagagtctattccgatcatcgacgtgtccaactgggacgaccccaaagtaggcgaggcgatttgcgacgcttcggagaaatggggtttctttcagatcatcaaccatgacatccccctggaagtgcttgacaacgttaaaaaggcgacatatcgcttctttgacttgtccgcagaggagaagaacaagtattccaaagagaataccgtgtccaacaatgcccgatacaccaccagctttattcctacggttgagaaggctctagaatggaaagattacctgagtctcttctatgtttccgaggaagaggcagctgctatctggccttcctcttgcaaggacgaagtgttagcttacatccataacggcgaaaaagttgcgaagaacttgttaaaggtgctactaaaggggctaaatgtgactgatatagacagtgagaaagagcaccttgtaacaggttcaaagagagttaacatgaacttctatcctaagtgcccaaaccctgatctcactgttggagttggtcgccattctgatgtctcggcatttactattctccttcaagatgatattggtgggctttacgtgagggtagagggtacggacagttgggttcatgtgccgccagtaaaagggtctcttgttattaatataggagatgctgtgcaaatacttagcaatggacgttacagaagtattgagcacagagtcgcagctaatggaaacagcgacaggatctcagttccccttttcatcaacccaaggccagaagatatcatcgctccgttacctgaagttctcgctaataccggagagaaaccgatttacaagccaattttgtactcggactactgcaggcacttttacaggaaggcccacgatggaaaggctaccattgatatcgcaaaggcataa

>FWZ796_39777967_39777967_C2’H4_Mal_HE_154X_1_chr005g000370

atggcgcctaccaaaaccagtgccgcacaatgggataacgattctgtaactgagtgggtgataaacaaaggcaatggagtgaagggtctctcagaaactggcctcaaaaccctcccccatcagtatatccagcccgtggaagagaggacgatgagtaaggtcttggtcggagagtctattccgatcatcgacgtgtccaactgggacgaccccaaagtaggcgaggcgatttgcgaagcttcagagaaatggggtttctttcagatcatcaaccatgacatccccctggaagtgcttgacaacgttaaaaaggcgacatatcgcttctttgacttgtccgctgaggagaagaacaagtattccaaagagaatactgtgtccaacaatgcccgatacaccaccagctttattcctacggttgagaaggctctagaatggaaagattacctgagtctcttctatgtttccgaggaagaggcagctgctctctggccttcttcttgcaagtaagactagtcgaaatctagctatttatttttaattaacattctcatacttacatgtaaatacacattataagttctatctctttcagattgccatgaccacaagtcctctgtatatataacaagtcatttcccaatgcataatacctacttattcatatattttaccatatatacttatctatacgtatatattaatttttcagggatgaagtgttagcttacatccataacggcgaaaaagttgcgaagaacttgttaaaggtgctactcaaggggctagatgtgactgacatagacagtgagaaagagcaacttgtaacaggttcaaagagggttaacatgaacttctatcctaagtgcccaaaccctgatctcactgttggagttggtcgccattctgatgtctcggcatttactattctccttcaagatgatattggtgggctttacgtgagggtagaggctacggacagttgggtccatgtgccgccagtaaaagggtctcttgttattaatataggtgatgctgtgcaaatacttagcaatggacgttacagaagtattgagcacagagtcgcagctaatggaaacagcgacaggatttcagttccccttttcatcaacccaagaccagaagatatcattgctccgttgcctgaagttctcgctaataccggagagaaaccgatttacaagccaattttgtactcggactactgcaggcacttttacaggaaggcccacgatggaaaggctaccattgatatcgcaaaggcataa

> Synthesized_C2’H5_Mal_HE_154X_1_chr005g002280

atggcgccttctaaacccaattgggccaacgattctgtcactgactttgtcataaccaaaggcaatggagtaaagggcctctcagaaactggcatcaaaacccttcccgaacaatatattcagcccctagaagagaggacaatgagtaaggtcatggtcggagagtctattccgatcatcgacgcgtctaactgggaggaccccaaagtcggggcggcgatatgcgatgctgcagagaaatggggtttctttcagatcatcaatcatggcataccccaggaagttcttgacaatgtcaaaaacgcgacatattgcttctttgactcgcctgctgaggagaaaaacaagtactccaaggagaattcggtatccaacaatgtccgatacaccacaagcttcattcctacagtcgagaaggctctggaatggaaagattacctcagtctcttctatgtttccgaggatgaggcctctaagttgtggccttcttcttgcagggatgaagtgttggcatacatgcataatggaagcatagttgtgaagcgactgttaagtgtgctactcaagggattaaatgtgactgaaatagacagtgaaaaagaacagcttgttacaggttcaaagagggttaacctcaatttctatcccatttgcccaaacccggaactcactgttggagttggccgccactctgatgtctcggcatttaccgttctccttcaagatgatatcggtgggctttatgtccgagtggagggtaaggacagttgggttcatgtgccgccggtaaaagggtctcttgttatcaacataggtgacgcattgcaaatatttagtaatgggcgttatagaagtattgagcacagagtcgtagctaatggaaatagtaacaggatctcagtccctcttttcattaacccaaggccagaggatatcattgctccattgcctgaagtgctcgctagtactggagagaaaccgatttacaaaccaattttgtactcggactacgtcaggcacttctacaggaagtcacacgatggaaagcttaccattgattttgcgaaggcataa
